# Atlas of Lysosomal Aging Reveals a Molecular Clock of Storage Disorder-Associated Metabolites

**DOI:** 10.1101/2025.09.25.678303

**Authors:** Anna M. Puszynska, Thao P. Nguyen, Andrew L. Cangelosi, Andrea Armani, Justin M. Roberts, Kristin A. Singh, James C. Cameron, Tenzin Tseyang, Grace Y. Liu, Steven Lai, Hans-Georg Sprenger, Jason Yang, William N. Colgan, Jibril F. Kedir, Kathrin M. Kajderowicz, Theodore K. Esantsi, Yuancheng Ryan Lu, Millenia Waite, Tenzin Kunchok, Caroline A. Lewis, Fabian Schulte, George W. Bell, David M. Sabatini, Jonathan S. Weissman

## Abstract

Lysosomal dysfunction is a well-recognized feature of aging, yet its systematic molecular investigation remains limited. Here, we employ a suite of tools for rapid lysosomal isolation to construct a multi-tissue atlas of the metabolite changes that murine lysosomes undergo during aging. Aged lysosomes in brain, heart, muscle and adipose accumulate glycerophosphodiesters and cystine, metabolites that are causally linked to juvenile lysosomal storage disorders like Batten disease. Levels of these metabolites increase linearly with age, preceding organismal decline. Caloric restriction, a lifespan-extending intervention, mitigates these changes in the heart but not the brain. Our findings link lysosomal storage disorders to aging-related dysfunction, uncover a metabolic lysosomal “aging clock,” and open avenues for the mechanistic investigation of how lysosomal functions deteriorate during aging and in age-associated diseases.

**One-Sentence Summary:** Aging in mice is tracked by a lysosomal “clock”, where glycerophosphodiesters and cystine – metabolites causally linked to juvenile lysosomal storage disorders – gradually accumulate in lysosomes of the brain, heart, skeletal muscle and adipose tissue.

## Introduction

Historically, much of aging research has focused on the pathobiology of age-related diseases (*1–7*), phenotypes associated with the gradual decline of organismal function (*8–12*), and the genetic dissection of signaling pathways that modulate lifespan in model organisms (*13–22*). In recent years, large-scale efforts employing single-cell transcriptomic approaches have provided an unprecedented tissue and cell-specific portrait of aging (*23–29*). Yet, transcript levels capture only one aspect of cellular physiology. Metabolites offer a complementary view of cellular biochemistry by providing a snapshot of the biochemical activities within tissues. Comprehensive studies on age-associated changes in metabolites have only recently begun to emerge (*30–33*), and those specifically interrogating individual organelles are missing.

From soon after the discovery of lysosomes (*34, 35*), the disruption of lysosomal functions has been hypothesized to be a central feature of the aging process (*10, 36–49*). Lysosomes are now recognized not only as a cellular waste disposal and recycling system (*36, 39, 50*), but also as regulators of cellular homeostasis, with complex functions in coordinating metabolism (*51–54*), responding to stress (*55–60*), and modulating signaling pathways that regulate lifespan, such as mTOR (*61–66*).

The importance of lysosomes to organismal health is underscored by the lysosomal storage disorders in which genetic mutations disrupt specific lysosomal enzymatic or transport activities (*67–72*). In this rare yet severe group of diseases, the accumulation of undegraded metabolite intermediates trapped within the lysosomal lumen leads to progressive cellular toxicity and organ damage (*73–75*). In recent years, compromised lysosomal function has been increasingly proposed as a contributing factor to age-associated neurological diseases, including Alzheimer’s disease (*76–79*) and Parkinson’s disease (*80, 81*), with a renewed appreciation that the aging lysosome may act as a “catalyst” of age-associated cellular decline (*82*).

Hence, elucidating the precise molecular nature by which the lysosome becomes compromised during aging and contributes to the aging process has become a requisite for developing new strategies to improve the healthspan and lifespan of the aging population. However, because lysosomes represent a small fraction of the cell volume, changes in their composition typically have a minimal impact on total cellular metabolite levels. Here, we used new and existing (*72*) tools for the rapid affinity purification of lysosomes to define, via unbiased mass spectrometry, their metabolite alterations during aging.

## Results

### Lysosomal *MeTabula Muris Senis* surveys the metabolome of the lysosome across aged tissues

To systematically characterize the metabolite composition of the aging lysosome, we used the recently developed LysoTag mouse line (*72*) (Fig. 1A), which harbors a lysosomal 3xHA-epitope tag that enables rapid lysosome isolation from any tissue of the body. We aged these mice to 24-26 months, a stage equivalent to a human in their 70s, and verified that constitutive expression of the LysoTag throughout the mouse lifespan does not appreciably alter the transcriptome or metabolome of tissues (Fig. S1). Apart from a few transcripts in LysoTag gastrocnemius muscle and metabolites in the aged LysoTag heart, LysoTag mice aged similarly to wildtype animals (Fig. S1), with no observable differences in morbidity or lifespan, suggesting that LysoTag mice accurately reflect the aging physiology of their wildtype counterparts. We next established efficient lysosome purification workflows from twelve tissues (Fig. 1A): brain, gut (small intestine), heart, kidney, liver, lung, pancreas, reproductive tissues (testis in males and uterus with the attached ovaries in females), skeletal muscle (gastrocnemius), spleen and white adipose tissue (perigonadal depot). In all analyzed tissues, the purity and yield of lysosomes were comparable between young and aged mice (Fig. 1B and S2).

**Fig. 1.**
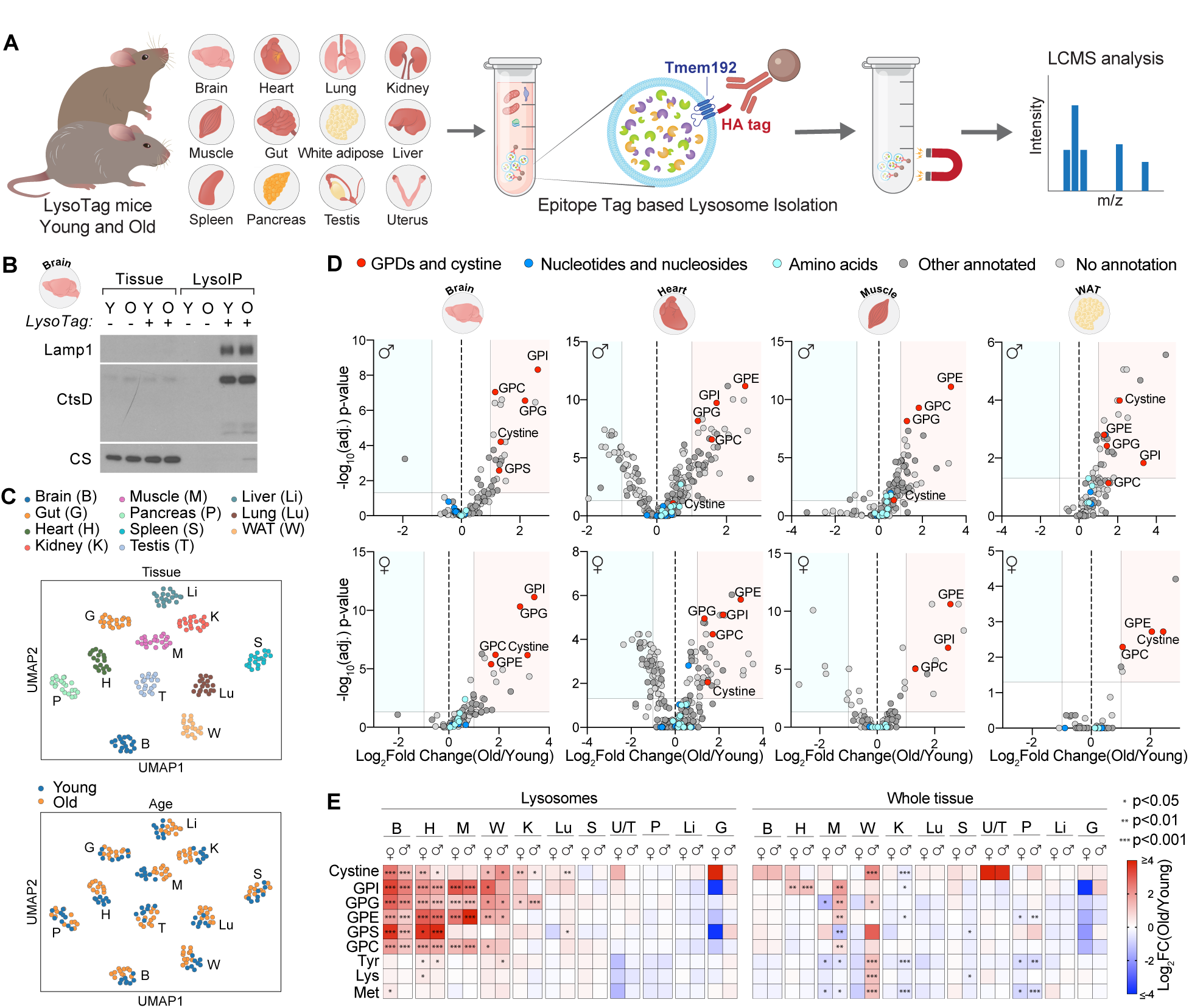
Atlas of lysosomal changes during aging reveals accumulation of cystine and glycerophosphodiesters in a subset of aged mouse tissues. (A) Schematic of the experimental setup for profiling the *in vivo* lysosomal polar metabolome. Lysosomes were isolated from tissues of young (2-3 months old) and old (24-26 months old) LysoTag mice using rapid immunoisolation with anti-HA magnetic beads. Global polar metabolomes were profiled using liquid chromatography coupled to high resolution mass spectrometry. (B) Levels of lysosomal and mitochondrial markers in tissue lysates and lysosomal fractions from the brains of young and aged LysoTag mice. Mock purifications from wildtype mice were performed to estimate nonspecific background. A representative sample from each group is shown. Full panels of organellar markers across all analyzed tissues are in Figure S2. Lamp1 – Lysosomal-associated membrane protein 1, CtsD – cathepsin D, marker of the lysosomal lumen; CS – citrate synthase, mitochondrial marker. (C) UMAP plots of lysosomal polar metabolomes from young (n = 8 to 9) and old (n = 11) male mice, colored by tissue of origin (top), or age group (bottom). Each data point represents a lysosomal fraction isolated from a specific tissue, with multiple tissues collected from the same mouse. Two separate cohorts of mice were used to cover all tissues represented in the UMAP. (D) Volcano plots showing untargeted lysosomal polar metabolomes from tissues of young (2-3 months old; n = 8 to 9) and old (24-26 months old; n = 11 to 12) male (top panels) and female (bottom panels) mice. The gray horizontal line indicates an FDR-corrected p-value = 0.05, the vertical lines represent log_2_(Fold change) of -1 and 1. GPC – glycerophosphocholine, GPE– glycerophosphoethanolamine, GPG – glycerophosphoglycerol, GPI – glycerophosphoinositol, GPS – glycerophosphoserine. (E) Heatmaps illustrating the log_2_-fold changes in lysosomal and whole tissue levels of selected metabolites between young and old mice, analyzed using a targeted method across all examined tissues. The dataset includes young (2-3 months old; n = 8 to 9) and old male (24-26 months old; n = 11) mice, as well as young (2-3 months old; n = 5 to 8) and old female mice (24-26 months old; n = 6 to 12). Asterisks denote FDR-corrected p-values (* p < 0.05, ** p < 0.01, *** p < 0.001). Full panel of analyzed metabolites is presented in Figure S5. Tissue abbreviations: H – Heart, B – Brain, M – Muscle (gastrocnemius), W – White adipose tissue (perigonadal depot), K – Kidney, Lu – Lung, S – Spleen, U/T – Uterus/Testis, P – Pancreas, Li – Liver, G – Gut (small intestine).

To construct a comprehensive atlas of how lysosomal metabolites change during aging – the Lysosomal *MeTabula Muris Senis* – we profiled the polar metabolomes of lysosomes isolated from young (2-3 months old) and aged (24-26 months old) male and female LysoTag mice. Unbiased clustering grouped the lysosomal metabolomes based on their tissue origin (Fig. 1C, top panel), indicating that different tissues exhibit unique metabolite landscapes within their lysosomes. Notably, young and aged lysosomes distinctly separated within most of the tissue-driven clusters (Fig. 1C, bottom panel), indicating widespread age-associated alterations in the metabolite composition of the lysosomes across tissues.

To identify candidate metabolites altered during aging, we performed in-depth untargeted analyses of lysosomes from each tissue (Fig. 1D, Fig. S3). While lysosomes from all analyzed tissues displayed distinct age-associated metabolite features, a common signature emerged across four tissues: aged lysosomes from the brain, heart, muscle, and white adipose tissue of both males and females contained highly elevated levels of cystine and metabolites belonging to the glycerophosphodiester (GPD) class, which are the end product of glycerophospholipid catabolism (Fig. 1D). Notably, GPDs and cystine accumulate in lysosomal storage disorders, with GPDs accumulating in Batten disease caused by *CLN3* mutations and cystine in cystinosis, as observed in patient samples (*68, 83–85*) and mouse models (*72, 84, 86*) of these diseases. Lysosomal accumulation of these molecules during healthy aging supports a long-hypothesized molecular link between lysosomal dysfunction of old age and juvenile lysosomal storage disorders. Our Lysosomal *MeTabula Muris Senis* reveals for the first time the identity of the molecular species involved.

We verified the annotations of GPDs and cystine with chemical standards (Fig. S4) and performed a targeted analysis of both the lysosomal and whole tissue fractions for high confidence quantifications (Fig. 1E and S5). This confirmed age-associated accumulation of GPDs and cystine within lysosomes of the brain, heart, muscle, and white adipose tissue, and revealed a partial phenotype in lysosomes of the kidney and male lung. Importantly, this lysosomal signature is not detectable at the whole tissue level, demonstrating the power of analyzing specific organelles to identify phenotypes that may be hidden or averaged out on a whole tissue level.

### Specific age-dependent accumulation of GPD and cystine is also a feature of rat aging

To enable studies of animals that are not readily genetically modifiable, we developed a “tag-free” approach for rapidly isolating lysosomes (Fig. 2A). We identified candidate antibodies that recognize cytosolic domains or loops accessible on four lysosomal proteins: p18 (LAMTOR1), cystinosin, Msfd12, and Tmem192. Among these, the anti-p18 antibody yielded the highest enrichment and purity of the isolated lysosomes based on organellar protein markers (Fig. S6A). Further, we validated that known lysosomal metabolites – amino acids and nucleotides (*54*) – are present in the “tag-free” lysosomes at levels above mock purifications, confirming the effective capture of intact lysosomes suitable for metabolomic analyses (Fig. S6B).

**Fig. 2.**
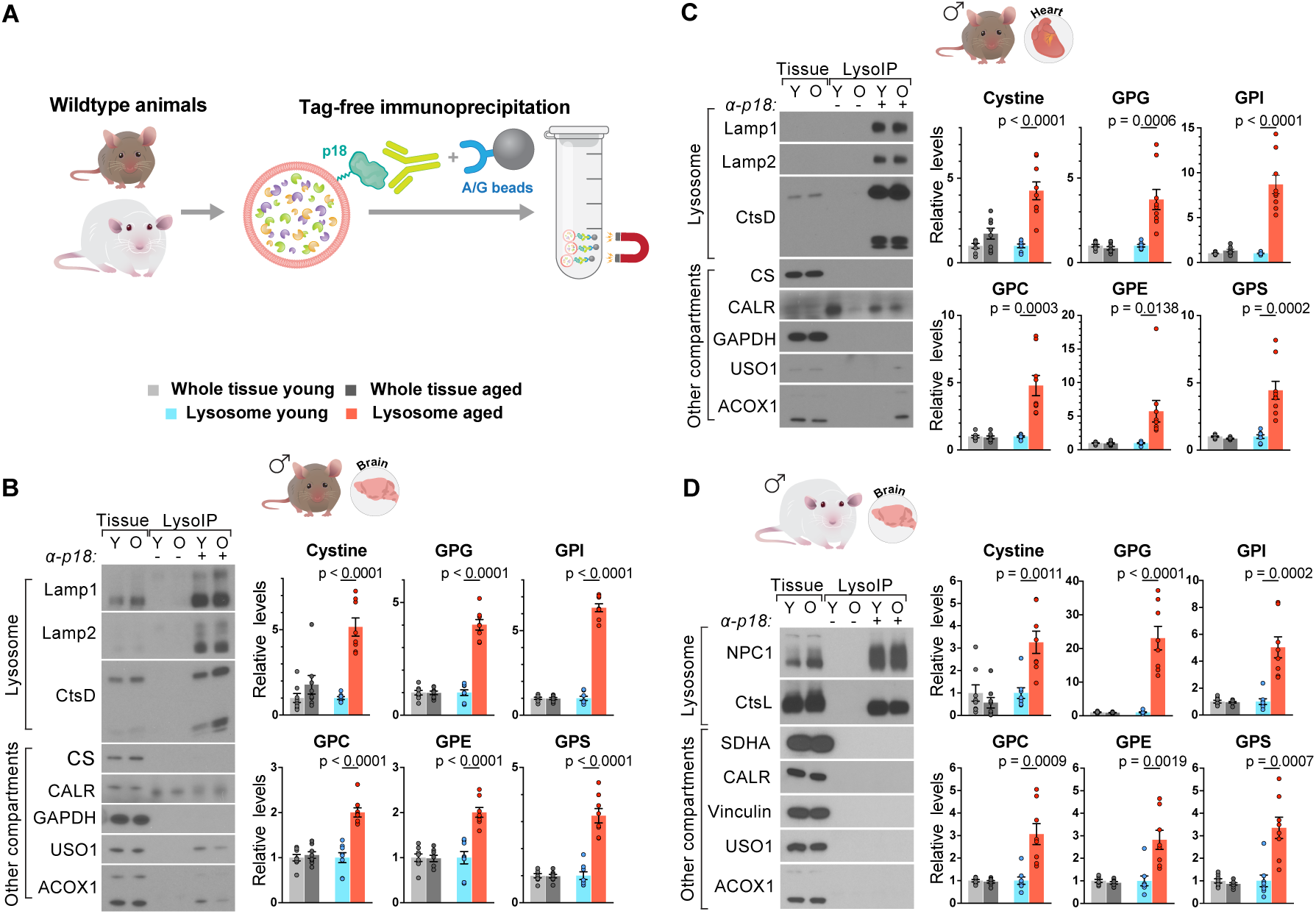
Lysosomes in aged wildtype mice and rats also accumulate cystine and glycerophosphodiesters. (A) Schematic of the experimental setup for the isolation of lysosomes from tissues of wildtype animals *in vivo* using an antibody against the lysosome-specific anchor protein p18. (B) Lysosomal fractions and whole tissue lysates from the brains of young (2-3 months old; n = 8) and old (24 months old; n = 8) wildtype mice were analyzed by immunoblotting for markers of lysosomes and other cellular compartments (left), and by targeted liquid chromatography and mass spectrometry for the quantification of cystine and glycerophosphodiesters (right). Lamp1 – Lysosomal-associated membrane protein 1, marker of the lysosomal membrane; Lamp2 – Lysosomal-associated membrane protein 2, marker of the lysosomal membrane; CtsD – cathepsin D, marker of the lysosomal lumen; CS – citrate synthase, mitochondrial marker; CALR – calreticulin, marker of the endoplasmic reticulum; GAPDH – Glyceraldehyde 3-phosphate dehydrogenase, cytoplasmic marker; USO1 – General vesicular transport factor p115, Golgi apparatus marker; ACOX1 – acyl-CoA oxidase 1, peroxisomal marker. (C) Lysosomal fractions and whole tissue lysates from the hearts of young (2-3 months old; n = 8) and old (24 months old; n = 9) wildtype mice were analyzed by immunoblotting for markers of lysosomes and other cellular compartments (left), and by targeted liquid chromatography and mass spectrometry for the quantification of cystine and glycerophosphodiesters (right). Markers probed as in (B). (D) Lysosomal fractions and whole tissue lysates from the brains of young (4 months old; n = 8) and old (25 months old; n = 8) wildtype rats were analyzed by immunoblotting for markers of lysosomes and other cellular compartments (left), and by targeted liquid chromatography and mass spectrometry for the quantification of cystine and glycerophosphodiesters (right). NPC1 – Lysosomal-associated membrane protein 1, marker of the lysosomal membrane; CtsL – cathepsin L, marker of the lysosomal lumen; SDHA – succinate dehydrogenase complex flavoprotein subunit A, mitochondrial marker; CALR – calreticulin, marker of the endoplasmic reticulum; Vinculin, cytoplasmic marker; USO1 – General vesicular transport factor p115, Golgi apparatus marker, ACOX1 – acyl-CoA oxidase 1, peroxisomal marker. (B-D) Data are presented as mean ± SEM. p-values were determined using two-tailed t-tests.

Using this “tag-free” strategy, we isolated lysosomes from the brain and heart of young (n = 8) and aged (n = 8-9) wildtype mice. Notably, the lysosomes from aged wildtype brain and heart had increased levels of GPDs and cystine similar to those of aged LysoTag mice (Fig. 2B and C), ruling out any potential confounding effects of the transgenic LysoTag on this age-associated lysosomal metabolite signature. Thus, lysosomal accumulation of GPDs and cystine in brain and heart is a *bona fide* signature of aging.

To determine whether other mammals also display this lysosomal aging signature, we first verified that our “tag-free” strategy can purify lysosomes from the tissues of rats, which are often considered a better model of human aging due to their metabolic similarities and longer lifespan. The protein and metabolite signature of isolated lysosomes confirmed the integrity of the rat immunoisolation protocol (Fig. S6C and D). We then obtained a cohort of young (4 months old; n = 8) and aged (25 months old; n = 8) rats from the National Institute on Aging. Lysosomes isolated from aged rat brains accumulated GPDs and cystine compared to those from the brains of young rats (Fig. 2D), establishing this lysosomal aging signature as common to rodent species. Inefficient mechanical disruption of the highly fibrotic aged rat hearts hindered the robust capture of lysosomes, preventing us from assessing lysosomes in this tissue. Despite this technical issue, we suggest that the lysosomal accumulation of GPDs and cystine in the brain – and likely in other tissues including the heart – may represent a newly-identified conserved hallmark of mammalian aging.

### Aged lysosomes accumulate bis(monoacylglycero)phosphate

To complement our analysis of the lysosomal polar metabolome, we also used the “tag-free” strategy to investigate compositional changes in the lysosomal lipidome during aging using unbiased untargeted lipidomics on lysosomes isolated from the brain, heart, muscle, and liver of young (3 months old; n = 7) and aged (24 months old; n = 7 to 9) wildtype male mice. A detailed lipid class analysis revealed that lysosomes from the brain, heart, and muscle – but not the liver – exhibit an age-associated increase in bis(monoacylglycero)phosphates (BMPs) and phosphatidylglycerols (PGs) (Fig. 3A). BMPs are unique lysosome-specific phospholipids (*87–90*) that serve as co-factors for lysosomal enzymes (*91, 92*), facilitating the degradation of other complex lipids, while PGs are their structural isomers (*87*). Increases in BMPs and BMPs/ PGs (representing isomeric BMPs or PGs that could not be distinguished) are particularly pronounced in brain and heart lysosomes, with fewer significantly elevated species in muscle (Fig. 3B). In contrast, the major lysosomal membrane phospholipids –phosphatidylcholines and phosphatidylethanolamines – remain largely unchanged in aged lysosomes. This is consistent with the notion that lysosomal yield was not increased in aged lysosome isolations and the increase in BMPs is compositional (Fig. 3A and B).

**Fig. 3.**
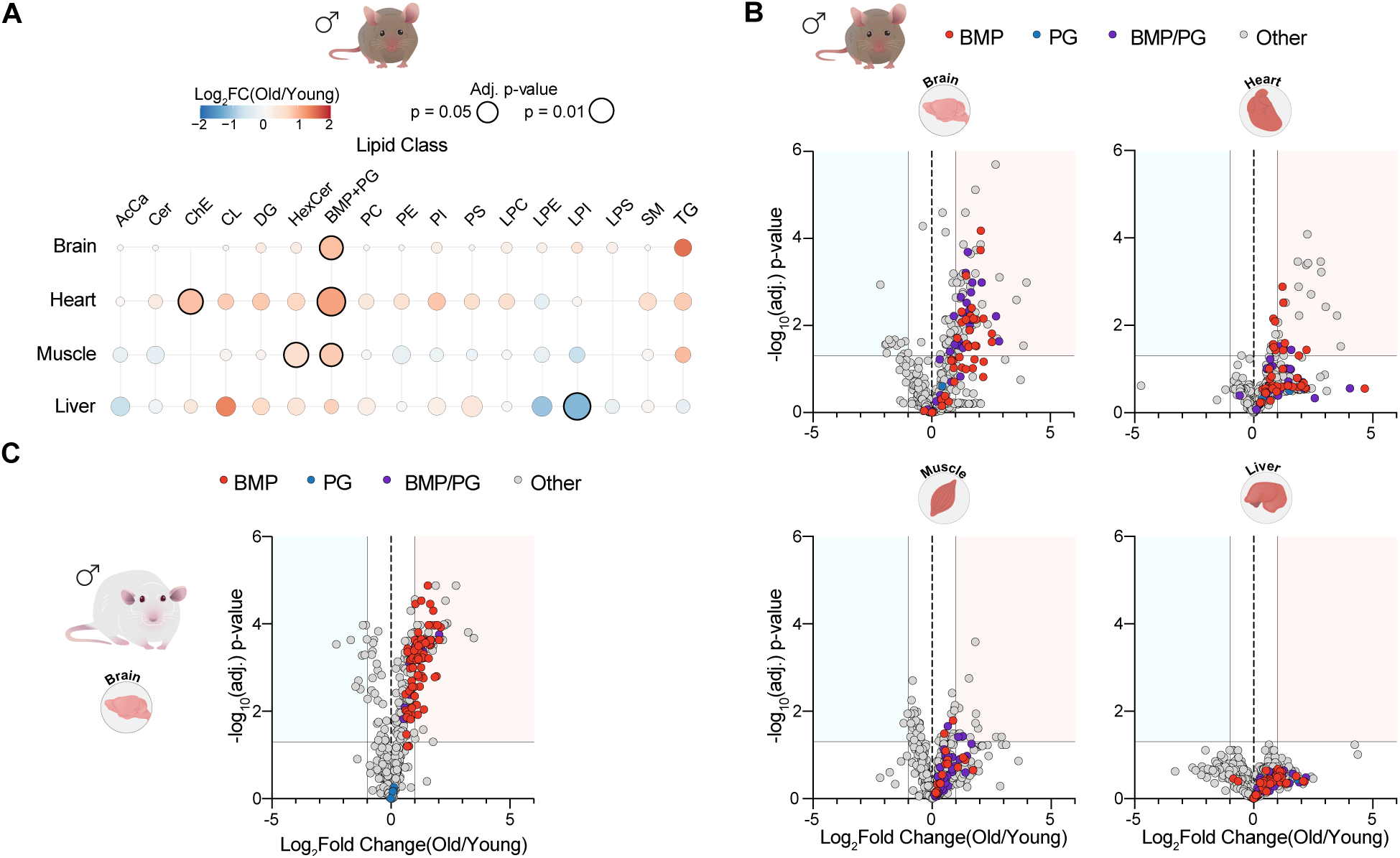
Lysosomes from aged mice show a compositional change in bis(monoacylglycero)phosphates. (A) Bubble plots representing differences in lipid classes in lysosomes purified from the brains, hearts, muscles, and livers of young (3 months old; n = 7) and old (24 months old; n = 7 to 9) wildtype mice. The color of each dot represents the log_2_-fold difference between old and young mice, while dot sizes are proportional to the FDR-corrected p-value. Lipid classes reaching FDR-corrected p < 0.05 are marked with black outlines. AcCa – Acylcarnitines, Cer – Ceramides, ChE – Cholesterol Esters, CL – Cardiolipins, DG – Diacylglycerols, HexCer – Hexosylceramides, BMP + PG – Bis(monoacylglycero)phosphates and Phosphatidylglycerols, LPC – Lysophosphatidylcholines, LPE – Lysophosphatidylethanolamines, LPI – Lysophosphatidylinositols, LPS – Lysophosphatidylserines, PC – Phosphatidylcholines, PE – Phosphatidylethanolamines, PI – Phosphatidylinositols, PS – Phosphatidylserines, SM – Sphingomyelins, TG – Triacylglycerols (B) Volcano plots of untargeted lysosomal lipidomes from the brains, hearts, muscles, and livers of young (3 months old; n = 7) and old (24 months old; n = 7 to 9) male mice. The gray horizontal line indicates an FDR-corrected p-value = 0.05, the vertical lines represent log_2_(Fold change) of -1 and 1. BMP species are highlighted in red, PG species are in blue, while BMP/PG species – representing isomeric BMPs or PGs that could not be distinguished – are in purple. (C) Volcano plot of untargeted lysosomal lipidomes from the brains of young (4 months old; n = 8) and old (25 months old; n = 8) male rats. The gray horizontal line indicates an FDR-corrected p-value = 0.05, the vertical lines represent log_2_(Fold change) of -1 and 1. BMP species are highlighted in red, PG species are in blue, while BMP/PG species – representing isomeric BMPs or PGs that could not be distinguished – are in purple.

A recent whole-tissue lipidomics study reported increased BMP levels in aged tissues (*93*); however, such analyses cannot distinguish between a true compositional shift within lysosomes and a general increase in lysosomal abundance. Here, we show that the BMP increase reflects a lysosome-intrinsic compositional change, paralleling the polar metabolite signature of elevated GPDs and cystine. Furthermore, we demonstrate that these lipid species are also elevated in lysosomes purified from the brains of aged rats (Fig. 3C). While BMP accumulation is a hallmark of certain lysosomal storage diseases (e.g., Niemann-Pick (*94*) and Gaucher diseases (*95*)), the aging-associated signature of elevated BMPs, GPDs, and cystine is not characteristic, to our knowledge, of any specific lysosomal storage disease and thus a unique feature of aged lysosomes.

Lysophospholipids are the immediate precursors of GPDs, both of which are generated during the process of phospholipid degradation by lysosomal phospholipases (*96*). In Batten disease, GPD accumulation has been linked to elevated levels of lysosomal lysophospholipids (*72, 86*). Increases in both glycerophosphocholine and lysophospholipids have also been associated with enhanced lipid trafficking to lysosomes (*97*), suggesting a coordinated accumulation driven by lysosomal phospholipid degradation.

In aged wildtype mice we observed no significant increases in any lysophospholipid class in the aged lysosomes from the brain, heart or muscle (Fig. S7A). We confirmed elevated levels of GPDs in these samples (Fig. S8), suggesting that, during aging, lysosomal GPD accumulation likely occurs independently of lysophospholipid storage. Similarly, in aged rat brain lysosomes, we observed increases in only a few specific lysophospholipid species, with no global elevation in this lipid class (Fig. S7B). As expected, given the absence of GPD elevation, aged mouse liver lysosomes also showed no increase in lysophospholipids (Fig. S7A and S8). When analyzing LysoTag mice, in addition to the increases in BMP levels, we observed an increase in lysosomal lysophospholipid levels (Fig. S9) in aged brain, heart and muscle, suggesting that these changes are impacted by the tagging system in this context.

### Microglia and neurons are the primary sites of lysosomal GPD and cystine accumulation in the aging brain

The brain is composed of a variety of cell types, including neurons, astrocytes, oligodendrocytes, and microglia, each with unique functions essential for maintaining brain homeostasis. These cell types not only vary in their physiological roles but also in how they respond to the aging process (*24, 26, 28*), highlighting the importance of understanding cell-specific contributions to age-related cognitive decline.

To investigate the lysosomal metabolome of specific cell types, we utilized Cre-driver mouse lines to express the LysoTag in four distinct brain cell populations – neurons, microglia, astrocytes, and oligodendrocytes – respectively achieved with Syn1-Cre, Cx3Cr1-Cre/ERT2, Aldh1l1-Cre/ERT2, and Plp1-Cre/ERT. LysoTag expression was faithfully restricted to each cell type in the respective mouse line (Fig. 4A) and efficiently induced by tamoxifen in young and aged mice of the tamoxifen-inducible lines (Fig. S10). We then optimized the lysosomal purifications from each of the four targeted cell types to attain robust enrichment of lysosomes from young and aged brains (Fig. 4B).

**Fig. 4.**
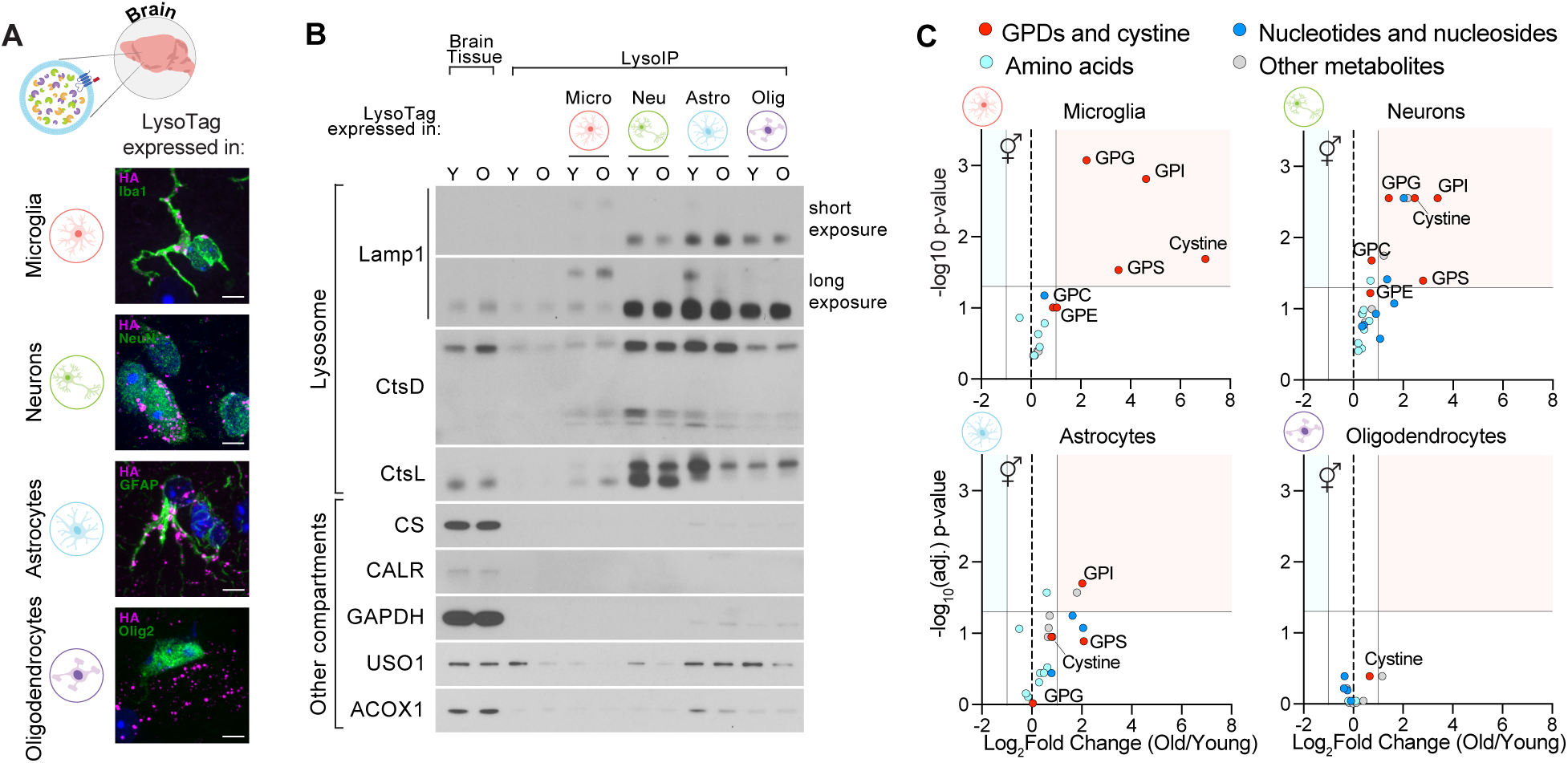
Lysosomes in microglia and neurons accumulate cystine and glycerophosphodiesters in the brains of aged mice. (A) Representative immunofluorescence images showing HA-tag expression (magenta) and cell-type specific markers (green) in brain sections from young mice (3 months old) expressing the LysoTag in specific cell types: microglia (Cx3Cr1-Cre/ERT2 LysoTag^flox/flox^ mice), neurons (Syn1-Cre LysoTag^flox/flox^ mice), astrocytes (Aldh1l1-Cre/ERT2 LysoTag^flox/flox^ mice) or oligodendrocytes (Plp1-Cre/ERT LysoTag^flox/flox^ mice). Iba1 (microglia), NeuN (neurons), GFAP (astrocytes), and Olig2 (oligodendrocytes) were used as markers. (B) Representative immunoblots of whole brain lysates and cell-type-specific lysosomal fractions from young (2-6 months old) and old (24-27 months old) mice, analyzed for markers of lysosomes and other cellular compartments. Markers probed as in Figure 2. (C) Volcano plots of cell type-specific lysosomal targeted polar metabolomes from the brains of young (2 to 6 months old; n = 5 to 8) and old (24 to 27 months old; n = 5 to 7) mice. Only metabolites significantly enriched above mock purifications are shown. The gray horizontal line indicates an FDR-corrected p-value = 0.05, the vertical lines represent log_2_(Fold change) of -1 and 1.

To assess potential biases in lysosomal enrichment across age groups, we performed quantitative TMT-based proteomics. This analysis confirmed comparable levels of total lysosomal proteins between young and aged mice, with only a modest (∼30%) increase observed in microglia, astrocytes, and oligodendrocytes from aged brains (Fig. S11A and B). Overall, lysosomal protein composition remained largely consistent across age groups, with few proteins showing significant differences between lysosomes purified from young and aged mice (Fig. S11B).

We then profiled the lysosomal polar metabolome of each brain cell type by targeted metabolomics (Fig. 4C). Notably, lysosomal GPDs and cystine were elevated in some, but not all, of the analyzed cell types, with aged microglial lysosomes displaying the starkest increase over young levels. Neuronal lysosomes also accumulated GPDs and cystine in the aged brain, establishing that this aging signature characterizes lysosomes of these specific brain cell types. As our proteomics analyses detected only modest differences in lysosomal protein yields from young and aged brains, this metabolite accumulation cannot be explained by variations in yield and primarily represents a genuine compositional change in the lysosomal metabolome of aged microglia and neurons. Conversely, lysosomes of astrocytes and oligodendrocytes exhibited minimal metabolite differences upon aging, with only glycerophosphoinositol significantly elevated in aged astrocytic lysosomes (Fig. 4C).

### Accumulation of GPD and cystine constitutes a metabolic “clock” of chronological and physiological aging

One of the key challenges in understanding age-related diseases is to distinguish primary aging phenotypes from secondary effects. Examining how age-associated traits correlate with chronological age can help identify intrinsic aging-related changes to prioritize as potential biomarkers or intervention targets. To this end, we quantified lysosomal levels of GPDs and cystine in wildtype male (n = 47) and female (n = 33) mice across their lifespans, from 77 to 985 days of age for males and 85 to 863 days for females. We focused on the brain and heart, where lysosomal accumulation of the GPDs and cystine is most pronounced with aging. In lysosomes from both tissues and sexes, isolated using our ‘tag-free’ strategy, cystine and GPD levels begin increasing early in life and continue to rise with age (Fig. 5A-B and S12), suggesting that they are an intrinsic feature of aging rather than merely a consequence of late-life organismal decline. Their accumulation follows a linear trend with age (Fig. 5A-B and S12), with r² values comparable to those reported for other aging clocks, including recently described spatial transcriptomic clocks that track aging in the brain (*29, 98–100*). Conversely, lysosomal levels of select amino acids and lysosomal protein yields were not appreciably altered with age (Fig. 5A-B, S12 and S13) indicating that this phenotype cannot be explained by differences in lysosomal yields. Notably, the chronological increase in lysosomal GPDs and cystine is not detectable at the whole cell level (Fig. 5A-B and S12) and thereby characterizes a previously unrecognized metabolic aging “clock” within lysosomes.

**Fig. 5.**
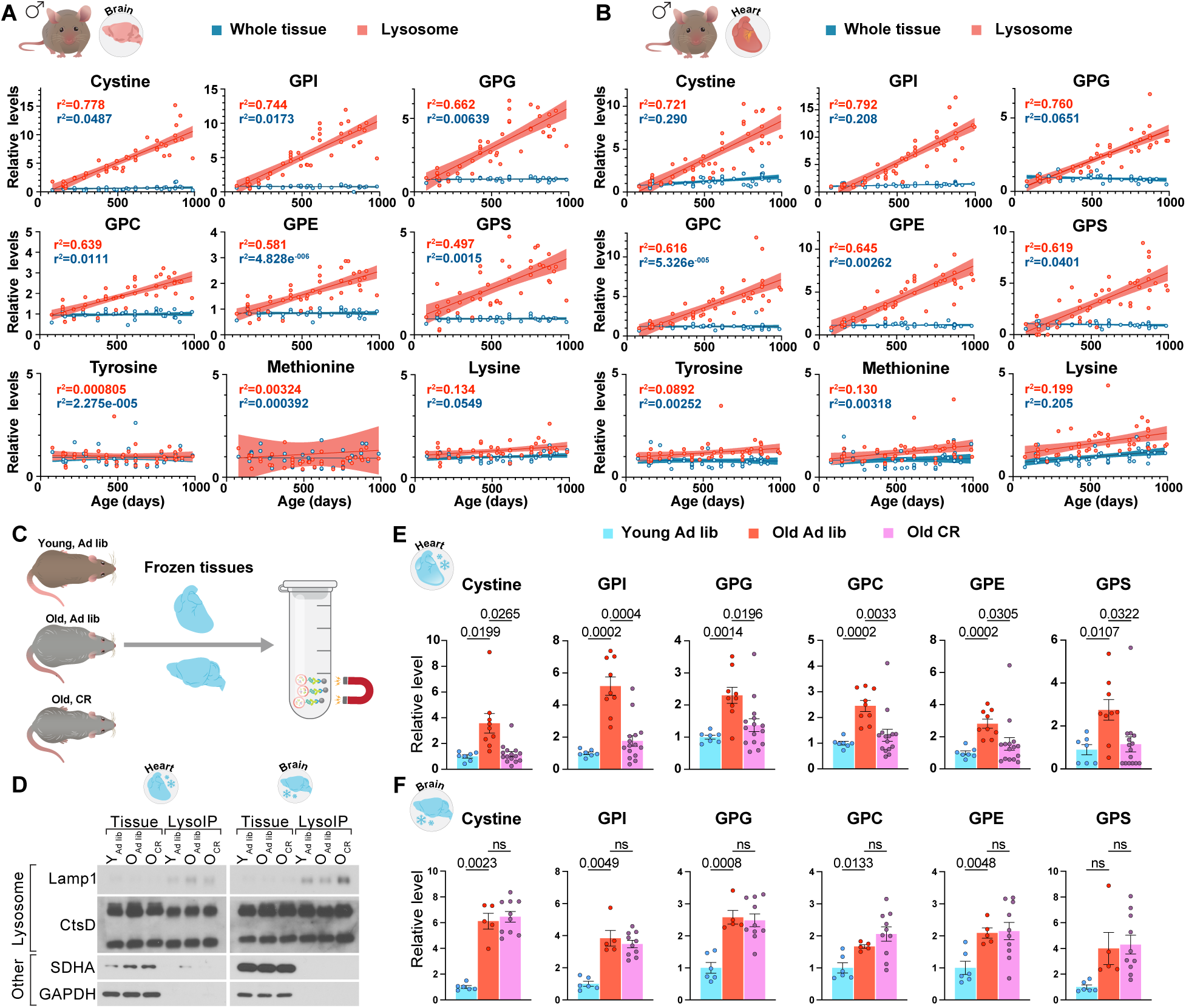
Lysosomal levels of cystine and glycerophosphodiesters increase with chronological age in wildtype mice and can be modulated by caloric restriction. (A) Levels of cystine, glycerophosphodiesters, and selected amino acids in brain lysosomes and whole tissue from male mice (n=47) ranging in age from 77 to 985 days. Shaded regions indicate 95% confidence intervals around the regression line. (B) Levels of cystine, glycerophosphodiesters, and selected amino acids in heart lysosomes and whole tissue from male mice (n=47) ranging in age from 77 to 985 days. Shaded regions indicate 95% confidence intervals around the regression line. (C) Schematic of the experimental setup for lysosome isolation from frozen tissues of wildtype animals. Ad lib – ad libitum fed, CR – caloric restricted. (D) Representative immunoblots of lysosomal fractions and whole tissue lysates from frozen hearts and brains of young (4 months old) and old (24 months old for heart, 28 months old for brain) wildtype mice, analyzed for markers of lysosomes and other cellular compartments. Markers probed as in Figure 2. (E) Lysosomal fractions from the hearts of young ad libitum-fed (4 months old; n = 7), old ad libitum-fed (24 months old; n = 10) and old caloric-restricted (24 months old; n = 15) wildtype male mice were analyzed by targeted liquid chromatography and mass spectrometry for quantification of cystine and glycerophosphodiesters. (F) Lysosomal fractions from the brains of young ad libitum-fed (4 months old; n = 6), old ad libitum-fed (28 months old; n = 5) and old caloric-restricted (28 months old; n = 10) wildtype male mice were analyzed by targeted liquid chromatography and mass spectrometry for quantification of cystine and glycerophosphodiesters. (E and F) Data are presented as mean ± SEM. p-values were determined using the Brown-Forsythe and Welch ANOVA test.

Next, we investigated if well-established anti-aging interventions could modulate this aging clock of the lysosome. In particular, caloric restriction is the most robust and well-documented intervention for delaying aging and improving metabolic health across a wide range of organisms, from yeast to rodents and non-human primates (*101–104*). To determine the impact of caloric restriction we analyzed frozen tissues of young and aged mice with lifelong caloric restriction or ad libitum feeding obtained from the NIA Tissue Bank. We first confirmed that our “tag-free” lysosome purifications work effectively with frozen tissue. Lysosomes purified from brain hemispheres flash-frozen and stored for six weeks retained detectable levels of GPDs and cystine compared to lysosomes from freshly-processed contralateral hemispheres, despite slightly lower and more variable yields (Fig. S14A-B). Further, the age-associated increase in lysosomal GPDs and cystine was detectable when comparing frozen young and aged brains (Fig. S14C), indicating that lysosomes isolated from frozen samples preserve biologically relevant metabolite contents.

We then purified lysosomes from frozen hearts and brains of young ad libitum-fed mice, aged ad libitum-fed mice, and aged caloric-restricted (CR) mice from the NIA Tissue Bank (Fig. 5C and D). In the heart, caloric restriction fully restored lysosomal cystine and GPDs to young levels (Fig. 5E). In contrast, caloric restriction had no detectable impact on cystine and GPD accumulation in lysosomes of the aged brain (Fig. 5F). Thus, this lysosomal aging signature is modulated by anti-aging interventions and reveals tissue-specific subcellular responses to such treatments, highlighting its potential as a measure of physiological age and as a tool to define the lysosomal basis of therapeutic strategies to counter aging.

## Discussion

Despite the long-standing notion that aging is associated with lysosomal dysfunction (*36–47*), a clear understanding of how lysosomes become altered during the aging process has remained elusive. Our study presents the first application of organellar metabolomics to aging – a Lysosomal *MeTabula Muris Senis* – providing a comprehensive portrait of the aging lysosome. We reveal tissue-specific patterns of lysosomal metabolite alterations that reflect both shared and unique features of aging across the mouse body. Critically, we define key compositional changes in polar metabolites of the aging lysosome, with GPDs and cystine accumulating with age in a subset of tissues. These alterations are independent of the LysoTag and are found in both mice and rats raised in distinct conditions, arguing that they represent a robust and conserved feature of the aging process. The increasing lysosomal load of GPDs and cystine progresses throughout an animal’s lifespan and emerges before its functional decline – mirroring the behavior of established aging clocks (*29, 98, 99*). This trend enables us to define a lysosomal aging metric, marking the first organelle-based biological clock and expanding the growing landscape of clocks built on metabolic (*33*) and molecular metrics (*29, 100*) beyond epigenetics. Notably, these lysosomal changes are not detectable at the whole-tissue level and coincide with compositional shifts in lysosomal BMPs. Finally, caloric restriction, a well-established anti-aging intervention, modulates GPDs and cystine levels in the heart but not in the brain. We speculate that the lack of a response to caloric restriction in brain lysosomes may be due to the nutrient homeostasis and metabolic adaptations in the brain that prioritize continuous energy supply during fasting.

One hypothesis that emerges from our work is that aging recapitulates phenotypes that occur in an accelerated manner in lysosomal storage disorders. While the connection between aging and these disorders has been hypothesized since the discovery of the lysosome (*36, 39, 40*), up to now there has been no insight into what metabolites are impacted. We establish phenotypic links between aging and Batten disease by identifying the shared accumulation of glycerophosphodiesters associated with lysosomal dysfunction caused by *CLN3* mutations (*72, 84–86*). We also draw connections to cystinosis, a disorder caused by defects in the lysosomal cystine transporter cystinosin, through the shared accumulation of cystine (*70, 83*). Finally, we observe compositional increases in the levels of BMPs known to be elevated in a subset of lysosomal storage disorders (*94, 95*). These parallels suggest that molecular pathways disrupted in lysosomal storage disorders may also be relevant to the metabolic dysregulation observed during aging, providing a foundation for exploring shared mechanistic underpinnings and potential common therapeutic targets across these conditions.

A critical question remains unanswered: are the accumulating polar metabolites and lipids merely byproducts of inefficient catabolism, or do they actively disrupt lysosomal or cellular functions? In the context of lysosomal storage disorders, accumulation of GPDs and cystine is closely linked to toxicity (*72, 86, 105, 106*) and leads to a functional decline and, eventually, premature death of the organism. We hypothesize that the buildup of these molecules in aged lysosomes could have deleterious effects on cells and tissues. Establishing the mechanisms of toxicity and evaluating whether interventions that specifically impact the levels of GPDs, cystine, and BMPs modulate aspects of the aging process may be critical for the development of future therapeutic approaches to promote healthy aging and extend lifespan.

Finally, lysosomal dysfunction is implicated in age-related neurodegenerative diseases such as Parkinson’s and Alzheimer’s. In these disorders, impaired lysosomal acidification (*107, 108*), defective lipid and protein degradation (*109*), and disrupted trafficking (*110*) have been suggested to contribute to the accumulation of neurotoxic aggregates such as α-synuclein and amyloid-β. A recent study further reinforces this link, highlighting ApoE4 as a contributing factor to lysosomal lipid dyshomeostasis (*111*). Understanding how ApoE4 and other genetic risk factors modulate the lysosomal aging “clock” – and whether the key metabolites we describe contribute to disease pathology – will be an important area for future research.

In conclusion, we provide a foundation for understanding lysosomal contributions to aging and establish a technical and conceptual framework for future investigations into the mechanisms driving these changes, their toxicity, and potential reversibility. Our findings highlight the lysosome as both a marker and likely mediator of aging. The ability to reset or slow the lysosomal aging “clock” presents an opportunity for therapeutic development, with the potential to mitigate not only lysosomal dysfunction but also its downstream effects on cellular health and aging phenotypes.

## Acknowledgments

We thank all members of the Weissman and the Sabatini lab, for suggestions and experimental help; We thank H. Lodish, T.M. Bertozzi and N. Sun for discussions and comments on the manuscript. We thank P.A. Sharp for advice. We thank Y. Ambaw, T. Walther and R. Farese for preliminary lipidomics analyses. We thank D. Valleau and S. Lourido for teaching us the S-Trap proteomics protocol. We are grateful to S. Sinha, R. Vallejo, B. Pham and C. Muresan for the technical and administrative support. We thank W. Chung for technical help, and B. Petrova for discussions and for sharing a spectral library. We thank the DCM veterinary team and the DCM Whitehead Mouse Facility for their support in maintaining the aging mouse cohorts. We thank the MIT BioMicroCenter for RNA library preparation and sequencing, as well as the Hope Babette Tang Histology Facility at the Koch Institute and S. Holder for histology support. We are grateful to C. Rausch from Warbler Creative for assistance with illustrations. We thank A. Salmon and the NIA Aged Rodent Tissue Bank for providing samples, and the NIA Aged Rodent Colonies for a cohort of young and aged rats.

## Author contributions

Conceptualization: A.M.P., J.S.W., D.M.S.

Formal analysis: A.M.P., W.N.C., G.W.B., J.S.W.

Funding acquisition: A.M.P., D.M.S., J.S.W.

Investigation: A.M.P., T.P.N., A.L.C., A.A., J.M.R., K.A.S., J.C.C., T.T., G.Y.L, S.L., H.-G.S., J.Y., J.F.K., K.M.K., T.K.E., Y.R.L., M.W., T.K., C.A.L., F.S.

Methodology: A.M.P., S.L., T.T., M.W., T.K., C.A.L., F.S., D.M.S., J.S.W.

Project administration: A.M.P., D.M.S., J.S.W. Supervision: D.M.S., J.S.W.

Visualization: A.M.P., W.N.C., J.S.W.

Writing – original draft: A.M.P., D.M.S, J.S.W.

Writing – review & editing: A.M.P., T.P.N., A.L.C., A.A., J.M.R., K.A.S., J.C.C., T.T., G.Y.L, S.L., H.-G.S., J.Y., W.N.C., J.F.K., K.M.K., T.K.E., Y.R.L., M.W., T.K., C.A.L., F.S., G.W.B., D.M.S., J.S.W.

## Data and materials availability

The RNA sequencing data is available under a GEO accession number GSE292311. Raw metabolomics and lipidomics data will be uploaded to Dryad. Other data generated are available from the corresponding authors on request.

## Supplementary Materials

## Materials and Methods

### Mice and Rats

All animal work was performed in accordance with the Massachusetts Institute of Technology (MIT) Committee on Animal Care. Mice were housed in the Whitehead Institute for Biomedical Research animal facility under the husbandry and veterinary care of the Department of Comparative Medicine. They were maintained on a 12-hour light/12-hour dark cycle at a constant temperature of 21°C and fed irradiated chow (LabDiet, 5P76). Mice were housed in groups, with a maximum of five mice per cage, and aggressive males were separated to prevent fighting. Where needed, mice were treated for non-life-threatening conditions, such as ulcerative dermatitis, as directed by the veterinary staff. Mice with a low body condition score were provided with gel food (Bio-Serv, NutraGel) and warm pads placed under the cage for thermal support, following veterinary staff recommendations.

For all experiments, sample sizes were not pre-determined. The researchers were not blinded to experimental groups. Aged animals were excluded from the study if, upon dissection, disseminated cancer or large tumors were found that could significantly alter cellular metabolism. Additionally, animals were removed from the study for humane reasons following veterinary assessment, based on the following clinical signs:

- Inability to eat or drink
- Severe lethargy, indicated by a lack of response when gently prodded with forceps
- Severe respiratory distress at rest, such as deep abdominal breathing or gasping
- Severe balance and/or gait disturbance
- Body Condition Score below 2-
- Loss of more than 20% of body weight within seven days
- Tumor burden greater than 2 cm
- Tumor ulceration

To achieve whole-body constitutive expression of the LysoTag, mice harboring lox-STOP-lox LysoTag (JAX strain #035401) were crossed with CMV-Cre mice (JAX strain #006054). CMV-Cre induces permanent recombination, generating a constitutively expressed LysoTag allele, and was subsequently removed in further crosses. The mice with whole-body LysoTag expression were backcrossed 10 times to C57BL/6J mice to achieve congenicity and were subsequently aged for 24–26 months. Genotyping was performed as previously described (*72*). Mice with heterozygous expression of the constitutive LysoTag allele were used.

To achieve cell-type-specific LysoTag expression in the brain, lox-STOP-lox LysoTag mice, backcrossed three times to C57BL/6J mice, were crossed with:

- Syn1-Cre mice (JAX strain #003966) for neuronal expression
- Cx3Cr1-Cre/ERT2 mice (JAX strain #020940) for inducible microglial expression
- Aldh1l1-Cre/ERT2 mice (JAX strain #031008) for inducible astrocytic expression
- Plp1-Cre/ERT mice (JAX strain #005975) for inducible oligodendrocytic expression

Genotyping of the Cre allele was performed using the following primers:

5′-GCCTGCATTACCGGTCGATGCAACGA-3′ and

5′-GTGGCAGATGGCGCGGCAACACCATT-3′.

All brain cell type-specific experiments used mice with homozygous expression of the LysoTag in the indicated cell type. For subsequent crosses involving Syn1-Cre LysoTag mice, Syn1-Cre-expressing females were crossed to males without Syn1-Cre to avoid germline recombination in the offspring (*112*). Additionally, offspring were genotyped as previously described (*72*) to confirm the absence of a recombined LysoTag allele.

In the inducible mouse lines (Cx3Cr1-Cre/ERT2 LysoTag, Aldh1l1-Cre/ERT2 LysoTag, and Plp1-Cre/ERT LysoTag mice), cell-type-specific LysoTag expression was induced by oral gavage of tamoxifen (200 mg/kg; MilliporeSigma, T5648) for five consecutive days. Mice were analyzed three weeks after the last tamoxifen dose. This time point was chosen to avoid potential acute effects of tamoxifen administration. Furthermore, in Cx3Cr1-Cre/ERT2 mice, it has been reported that labeled circulating monocytes are cleared within 2–3 weeks after Cre induction, while microglial labeling remains stable for a prolonged period (*113*).

Male young (4 months old) and old (25 months old) F344-CDF rats were obtained from National Institute on Aging Aged Rodent Colonies, housed at the Charles River Laboratories (Frederick, MD). The animals were provided ad libitum access to chow and water, and were housed in pairs when imported with a cage mate; otherwise, they were housed individually. All rats were maintained on a 12-hour light/dark cycle and were acclimated to the facility for fourteen days prior to sample collection.

Frozen mouse hearts and brains were obtained from the National Institute on Aging Aged Rodent Tissue Bank. We obtained frozen hearts from young (4-month-old) and old (24-month-old) animals, as well as frozen brains from young (4-month-old) and old (28-month-old) animals, based on sample availability. The tissues were collected from three experimental groups: young ad libitum (AL), aged ad libitum (AL), and aged calorie-restricted (CR) mice. All mice were initially fed the NIH-31 chow diet (AL) until weaning. At weaning, mice were separated into AL and CR groups. Mice in the AL group continued to receive the NIH-31 chow diet ad libitum, while CR mice were switched to the NIH-31 fortified diet. Caloric restriction (CR) was implemented progressively, beginning at 14 weeks of age with a 10% reduction in intake, followed by a 25% reduction at 15 weeks, and a 40% reduction at 16 weeks, which was maintained for the remainder of their lifespan. All CR animals were fed daily on a fixed schedule at approximately the same time each day, seven days a week. The NIH-31 fortified diet was used to prevent malnutrition as a consequence of CR (*114*). Water was provided ad libitum through an automated watering system. Tissue collection was performed under protocols approved by the appropriate institutional guidelines for animal care and use.

### Rapid LysoTag Lysosome Purification

For LysoTag immunoisolation from fresh mouse tissues, tissues were collected and processed immediately after euthanasia. For the brain, the cerebral hemispheres were dissected, and each was used for a separate isolation. For the liver, 130 mg of the major lobe was weighed. For the heart, the whole heart was excised, cleared of blood in PBS, minced on an ice-cold metal block with razor blades, and a set amount of minced tissue (80 mg for female tissues and 90 mg of male tissue) was weighed. For the muscle, the gastrocnemius muscle was dissected and separated from the soleus muscle, minced on an ice-cold metal block, and a set amount of minced tissue (80 mg for female tissues and 90 mg of male tissue) was weighed. For white adipose tissue, the perigonadal white adipose depot was excised and 130 mg was weighed. For the lung, the lung was excised, cut into smaller pieces, and 70 mg of tissue was weighed. For the pancreas, the pancreas was excised, and 100 mg of tissue was weighed. For the kidney, 110 mg of excised tissue was weighed. For the spleen, 50 mg of excised tissue was weighed. For the gut, a 5-cm section of the gut, starting at the gastroduodenal junction, was excised, the mesenteric tissue was cleaned away, and the digested food was washed out with KPBS (136 mM KCl, 10 mM KH₂PO₄, pH 7.25 in Optima LC–MS water) using a round-tip needle (FST, 18061-50) inserted into the gut lumen. The gut was then minced on an ice-cold metal block, and 200 mg of minced tissue was weighed. For the testis, both testes were used for each isolation. For the uterus, the uterus was excised, cleaned of perigonadal white adipose tissue and minced on an ice-cold metal block. For each tissue, wildtype mice were used in mock purifications to establish nonspecific background.

In whole-body LysoTag-expressing mice, multiple tissues were rapidly collected and processed simultaneously by a team of investigators. In experiments where the pancreas was used for lysosome isolation, the pancreas tissue was excised first to avoid autolysis. In experiments where lung was collected, lung tissue was collected before heart excision to prevent contamination of the lung tissue with blood. For each tissue, care was taken to weigh the same amount of input material for all mice within an experiment.

Tissue disruption was performed in 1 ml of ice-cold KPBS (with cOmplete EDTA-free protease inhibitor cocktail (Roche) and PhosSTOP phosphatase inhibitor cocktail (Roche) for protein analysis, and without protease and phosphatase inhibitors for metabolite and lipid analysis) using a 2 ml Dounce homogenizer. Soft tissues (brain, liver, kidney, pancreas, spleen, WAT, testis) were homogenized using a 2 ml borosilicate glass tissue grinder with a plain plunger (VWR, 89026-386 and 89026-398), while tougher tissues (heart, muscle, lung, uterus, gut) were homogenized using a 2 ml borosilicate glass tissue grinder with a ground-glass pestle (Kimble Kontes, 885500-0021). All tissues required 25 strokes for optimal homogenization, except WAT, which required 35 strokes.

For whole tissue lysate analysis, 10 μL of the homogenate was mixed with either 90 μL of 80% methanol for metabolite extraction, 90 μL of 100% isopropanol for lipid extraction, or 90 μL of Triton Lysis Buffer (50 mM HEPES, pH 7.4, 40 mM NaCl, 1% Triton X-100, 2 mM EDTA, and protease and phosphatase inhibitor cocktails) for protein analysis. The remaining tissue lysate was centrifuged at 1,000 g for 2 min at 4 °C. The supernatant was then incubated with 100 μL of pre-washed anti-HA magnetic beads (Pierce) on a rotating shaker at 4 °C. For the kidney, the supernatant required filtration through a 100 μm mini-Strainer (PluriSelect, 43-10100-76) before incubation with magnetic beads to avoid small tissue fragments. For WAT, care was taken to avoid the fat layer on the surface of the supernatant. For all tissues, care was taken to prevent disturbing the tissue pellet.

The beads were incubated with the supernatants for 5 min for polar metabolomics and immunoblotting, 7 min for lipidomics, and 10 min for protein analysis via proteomics. After incubation, the beads were washed three times with 1 mL of ice-cold KPBS by placing the tubes on a DynaMag-2 magnet (Invitrogen, 12321D) to trap the beads and remove the supernatant. To increase purity, the sample was transferred to a new tube after the first and third washes. After the final resuspension, the bead suspension was split: 800 μL was used for metabolite or lipid extraction, while 200 μL was used for quality assessment of lysosomal isolation by protein quantification and immunoblotting.

The lysosomal fractions were resuspended in:

- 45 μL of ice-cold 80% methanol in LC–MS water containing 500 nM isotope-labeled amino acids (Cambridge Isotope Laboratories) for polar metabolite analysis.
- 45 μL of ice-cold isopropanol containing 1000 ng/ml SPLASH LIPIDOMIX internal standard mix (Avanti, 330707) for lipid analysis.
- 50 μL of Triton Lysis Buffer with protease and phosphatase inhibitors for protein analysis.

The input and lysosomal fractions for metabolite and lipid analysis were vortexed for 10 min at 4 °C and centrifuged at 21,000 g. For the input, the supernatant was collected. For the lysosomal fraction, tubes were placed on the magnet, and the eluate was transferred to a new tube and stored at -80 °C until analysis.

### “Tag-Free” Lysosome Isolation

The Tag-free lysosome isolations followed the same protocol with modifications. Briefly, postnuclear supernatants from each tissue were incubated with 2.5 μg of anti-p18 antibody (CST, 8975S; RRID: AB_10860252) for 7.5 min at 4 °C. Isotype-specific IgG (CST, 3900S; RRID: AB_1550038) was used in mock purifications to establish nonspecific background. 100 μL of pre-washed protein A/G magnetic beads were then added, followed by an additional 7.5 min incubation at 4 °C. Washes and extractions were performed as described above. In our initial protocol development phase, we also tested anti-CTNS (Biorbyt, orb247656), anti-Tmem192 (Huabio, HA721106; RRID: AB_3072230) and anti-MFSD12 (St John’s Laboratory, STJ195920) in both mouse and rat experiments.

For experiments performed using rat brain tissue, freshly harvested brain was separated into hemispheres, and one hemisphere was used for lysosome isolation. The hemisphere was disrupted in 4 ml of ice-cold KPBS without protease and phosphatase inhibitors using a 10 ml Dounce homogenizer with smooth PTFE pestle (Cole-Palmer, EW-44468-15 and EW-44468-03). Homogenization was performed with 30 strokes on ice. Each of three 2 mL tubes received 1.6 mL of homogenate for metabolomics, lipidomics, and mock lysosome immunoisolation. The homogenates were precleared by centrifugation at 1,000 g for 2 minutes at 4°C, and the resulting supernatant was incubated with 3 μg of either anti-p18 antibody (CST, 8975S) or an isotype-specific IgG control (CST, 3900S). The subsequent steps were carried out as described above.

For lysosome isolation from the frozen mouse tissue, the entire brain was used, except for the experiment presented in Figures S14 A and B, where one hemisphere was used fresh and the other frozen for comparison. The frozen brain was placed on an ice-cold metal block for 30 sec to soften, then transferred to a 2 ml Dounce homogenizer (VWR, 89026-386 and 89026-398) containing 1.5 ml of ice-cold KPBS. The entire homogenate was transferred to a 2 ml tube, and the subsequent steps were identical to those used for a fresh mouse brain. For the frozen hearts, each heart was placed on an ice-cold metal block and trimmed with a razor blade to obtain 160-180 mg of tissue for each lysosome isolation. The trimmed heart was transferred to a 2 ml Dounce homogenizer (Kimble Kontes, 885500-0021) without additional mincing, and homogenized with 20 strokes on ice. The following steps were performed as described above.

### Plasma collection for Metabolite Profiling

Blood was collected from the submandibular vein into lithium heparin-coated tubes (Sarstedt, 20.1292.100), followed by plasma extraction according to the manufacturer’s protocol. Plasma was diluted 1:9 (v/v) with an extraction buffer (75:25:0.2 acetonitrile:methanol:formic acid) containing isotopically labeled internal standards. The samples were vortex-mixed for 10 min, and centrifuged at 15,000 rpm for 10 min at 4°C to pellet debris. The resulting supernatant was collected for metabolite profiling.

### Polar Metabolomics

Metabolite profiling was performed using a QExactive bench-top Orbitrap mass spectrometer (Thermo Fisher Scientific, San Jose, CA) equipped with an Ion Max source and a HESI II probe, coupled to a Dionex UltiMate 3000 HPLC system. External mass calibration was conducted every 7 days using a standard calibration mixture. Typically, 5 μL of the lysosome sample and 2.5 μL of the whole tissue extract were injected onto a SeQuant® ZIC®-pHILIC 150 × 2.1 mm analytical column (5 μm particle size, EMD Millipore), fitted with a 2.1 × 20 mm guard column of the same material. The mobile phase consisted of Buffer A (20 mM ammonium carbonate, 0.1% ammonium hydroxide) and Buffer B (acetonitrile). The column oven was maintained at 25°C, and the autosampler tray at 4°C. Chromatographic separation was performed at a flow rate of 0.150 mL/min using the following gradient: 0–20 min: Linear gradient from 80% to 20% B, 20–20.5 min: Linear gradient from 20% to 80% B, 0.5–28 min: Hold at 80% B. The mass spectrometer operated in full-scan, polarity-switching mode, with the following parameters: spray voltage: 3.0 kV, heated capillary temperature: 275°C, HESI probe temperature: 350°C, sheath gas flow: 40 units, auxiliary gas flow: 15 units, sweep gas flow: 1 unit, mass range: m/z 70–1000, resolution: 70,000, AGC target: 1 × 10⁶, maximum injection time: 20 ms. For experiments in Figures 2, 5, S8, S12, S14, samples were run in positive mode only with a targeted selected ion monitoring scan (tSIM) centered on m/z *260.05298, 216.06320, 258.11010, 335.07380, 247.05773, 241.03112, 147.11280, 150.05830, 182.08120* for more accurate quantitation of GPS, GPE, GPC, GPI, GPG, Cystine, Lysine, Methionine and Tyrosine respectively. Data presented in S8 were acquired using the isopropanolic extracts prepared for lipidomics analysis.

For untargeted metabolomics data were acquired with data-dependent MS2 (ddMS2) analysis performed on pooled samples using a Top-10 method with stepped collision energies of 15, 30, and 45 V. The MS2 acquisition parameters were: resolution: 17,500, AGC target: 2 × 10⁵, maximum injection time (max IT): 100 ms, isolation window: 1.0 m/z. Data were processed using Compound Discoverer 3.3 (Thermo Fisher Scientific), incorporating an in-house mass list for metabolite identification and in-house spectral libraries. For the UMAP analysis, all male samples were aligned and processed together using Compound Discoverer 3.3. Total ion counts per sample were normalized to a sum of 10,000, followed by log transformation and z-scoring of metabolite intensities. UMAP was performed using the first 50 principal components, with 10 neighbors, a minimum distance of 2, and a spread of 2. For the volcano plots in Figures 1 and S3, each tissue was analyzed individually in Compound Discoverer 3.3.

For targeted metabolomics, relative quantification of polar metabolites was performed using TraceFinder™ 4.1 (Thermo Fisher Scientific), applying a 5 ppm mass tolerance and referencing an in-house library of chemical standards. Missing metabolite values were imputed using half of the minimum value of that metabolite across samples with values. Metabolite levels in a tissue were summarized by mean, and the confidence about any difference between old and young tissues was calculated with two-sided t-tests (with the Welch modification), followed by multiple-hypothesis correction with False Discovery Rate. Both lysosome and whole tissue lysate samples were analyzed in the same manner. For lysosomal analyses, metabolites were filtered out from the analysis if their level in both the young and old samples were less than 2-fold higher than the mock immunoisolation samples.

### Validation of glycerophosphodiesters and cystine annotation

LC-MS-based analyses of chemical standards and brain extracts were conducted on a Xevo™ G3 QTof benchtop mass spectrometer and a Select Series™ Cyclic IMS mass spectrometer. Both were coupled to an ACQUITY™ Premier UPLC liquid chromatography system with a quaternary solvent manager and a flow through needle autosampler (Waters). Liquid chromatography method was as described above.

The Xevo G3 QTof mass spectrometer was operated in electrospray positive and electrospray negative ionization mode, with a capillary voltage of 1.0 kV in positive ion mode and 2.0 kV in negative ion mode. The desolvation temperature was 450 °C and a nitrogen desolvation gas and cone gas flow of 1000 L/hr and 50 L/hr, respectively. The source operating temperature was 120 °C. Data was acquired using a Data Dependent Acquisition method, with the survey scan Cone Voltage at 40 V and the MSMS collision energy at 15 eV. Argon was used as the collision gas. The top 5 ions were selected for MSMS acquisition with a trigger threshold of 200 counts per second and an acquisition threshold of the total ion chromatogram (TIC) falling below 100 counts per second or a 1 second total acquisition time. Deisotope peak selection was turned off and no dynamic exclusion list applied. An inclusion list for Cystine, GPC, GPE, GPG, GPI and GPS was applied.

The Select Series Cyclic IMS mass spectrometer was used to confirm the identification of GPS in the extract samples. It was operated in electrospray negative ionization mode, with a capillary voltage of 2.0 kV. The desolvation temperature was 450 °C and a nitrogen desolvation gas and cone gas flow of 1000 L/hr and 50 L/hr, respectively. The source operating temperature was 120 °C. Data was acquired using an Ion Mobility Separation (IMS)-MSMS method, with the quadrupole selection window set to Unit Resolution (∼0.7 Da FWHM) for m/z 258. The cyclic Ion Mobility (cIM) cell used nitrogen as the drift gas, with a traveling wave height ramp of 12 to 16 V. The cone voltage was 10 V and the transfer collision energy set to 21 eV. Nitrogen was used as the collision gas.

### Preparation of GPG and GPS standards

Because commercial standards for GPG and GPS are not readily available, they were prepared in-house via hydrolysis of 14:0 lysophosphatidylglycerol (Avanti, 858120P) and 18:0 lysophosphatidylserine (Avanti, 858144P) respectively, following the standard alkaline hydrolysis reaction for phospholipids (*115*). Each lysophospholipid was hydrolyzed under alkaline conditions to generate glycerophosphodiesters. Briefly, 0.5 mL of a 0.132 M NaOH and 0.05 M KCl solution was added to 0.5 mL of a lysophospholipid dispersion in a glass vial, resulting in a final lysophospholipid concentration of 5 mM. The mixture was stirred at 40 °C for 1 hour. Following hydrolysis, the solution was neutralized with 6 M HCl. Glycerophosphodiesters were extracted using a standard biphasic chloroform/methanol/water extraction. The aqueous phase was collected, dried and resuspended in 80% methanol.

### Lipidomics

Lipid separation of the fractions collected from mouse tissues was performed using an Ascentis Express C18 column (2.1 × 150 mm, 2.7 μm; Sigma-Aldrich) on a Vanquish Horizon UPLC system coupled to an Orbitrap Exploris 480 mass spectrometer (Thermo Fisher Scientific) equipped with a heated electrospray ionization (HESI) probe. For fractions collected from rat brain tissue, lipid separation was carried out using an Accucore C30 column (2.1 × 150 mm, 2.6 μm; Thermo Fisher Scientific) on a Vanquish Horizon UPLC system coupled to an ID-X Tribrid mass spectrometer (Thermo Fisher Scientific). 4 μL of the isopropanol-extracted lysosome sample from the liver, or 5 μL from the brain, heart and muscle was injected onto the column, with separate injections for positive and negative ionization modes. The mobile phase composition was: Buffer A: 60:40 (v/v) water:acetonitrile with 10 mM ammonium formate and 0.1% formic acid; Buffer B: 90:10 (v/v) isopropanol:acetonitrile with 10 mM ammonium formate and 0.1% formic acid. Chromatographic separation followed a 45-minute gradient, adapted from Bird et al. (2011) (*116*): 0–1.5 min: Isocratic at 32% B, 1.5–4 min: Increase to 45% B, 4–5 min: Increase to 52% B, 5–8 min: Increase to 58% B, 8–11 min: Increase to 66% B, 11–14 min: Increase to 70% B, 14–18 min: Increase to 75% B, 18–21 min: Increase to 97% B, 21– 25 min: Hold at 97% B, 25–25.1 min: Reduce to 32% B, followed by 19.9 min of column re-equilibration. The flow rate was 0.260 mL/min. The column oven and autosampler were maintained at 55 °C and 15 °C, respectively. The mass spectrometer was operated in full-scan data-dependent MS/MS (ddMS2) mode, with the following parameters: MS1 Orbitrap resolution: 120,000; MS/MS Orbitrap resolution: 30,000; Spray voltage: 3.25 kV (positive mode) or 3.0 kV (negative mode); Heated capillary temperature: 300 °C; HESI temperature: 375 °C; S-lens RF level: 45; Sheath gas flow: 40 units; Auxiliary gas flow: 10 units; Mass scan range (m/z): 200–2000; AGC target: 1 × 10⁵; Maximum injection time: 50 ms. The following parameters were used for data-dependent MS/MS: Cycle time: 1.5 sec; Isolation window: 1.0 m/z; Intensity threshold: 1 × 10³; HCD fragmentation: Stepped collision energy at 15, 25, and 35 units; MS/MS AGC target: 5 × 10⁴; MS/MS maximum injection time: 54 ms; Isotopic exclusion: Enabled; Dynamic exclusion window: 2.5 sec. Internal calibration using Easy IC was enabled. Quadrupole isolation was applied for precursor selection.

For targeted metabolomics in Figure S9, relative quantification of lysophospholipids was performed using TraceFinder™ 4.1 (Thermo Fisher Scientific), applying a 5 ppm mass tolerance and referencing an in-house library of lipid species.

### Lipid Annotation and Quantification

High-throughput lipid annotation and relative quantification were performed using LipidSearch v5.0 (Thermo Fisher Scientific/Mitsui Knowledge Industries) (*117*) utilizing the HCD database. LipidSearch matched MS/MS spectra to the database with the following parameters: Precursor ion tolerance: 5 ppm, Product ion tolerance: 10 ppm. LipidSearch nomenclature follows a convention using underscores to denote the absence of sn-positional information (e.g., PC(18:0_18:1) rather than PC(18:0/18:1)). If insufficient MS/MS data were available to resolve all fatty acyl chains, only the sum composition was reported (e.g., PC(36:1)). After peak detection, positive and negative mode data were aligned with retention time tolerance 0.25 min. Each aligned peak was carefully inspected, and manual integration was performed when necessary to improve accuracy. Peaks identified as background were excluded from further analysis. BMP and PG species were annotated as described in (*72*). Briefly, the two lipid classes were manually annotated using MS2 fragmentation data acquired in positive ion mode. In cases where MS2 data were unavailable, the species was annotated as BMP/PG.

Raw peak areas for all annotated lipids were exported to Microsoft Excel, where data were filtered to retain only lysosomal peaks that were at least twofold higher than those in the mock purification. For lipid class analysis, the total area of each class was calculated by summing the areas of all lipid species within that class for each sample. BMP and PG species, as well as lipid features that could not be unequivocally assigned as either BMP or PG (BMP/PG), were grouped together and analyzed as a single class (denoted BMP+PG). To ensure consistency in sample loading, isotope-labeled internal standard lipids were assessed in each sample to verify uniform injection volumes and the absence of signal variations.

### Proteomics

Protein lysates were prepared in Triton Lysis Buffer (50 mM HEPES, pH 7.4, 40 mM NaCl, 1% Triton X-100, 2 mM EDTA, and protease and phosphatase inhibitor cocktails) and processed for proteomics using S-Trap columns (Protifi). Briefly, SDS was added to a final concentration of 5%. Proteins were reduced with 5 mM tris(2-carboxyethyl)phosphine (TCEP; Pierce, 20490) for 10 min at 55 °C, then alkylated with 20 mM methyl methanethiosulfonate (MMTS; Pierce, 23011) for 10 min at room temperature. Next, samples were acidified with phosphoric acid to a final concentration of 2.5% and the proteins were precipitated using S-Trap binding buffer (90% methanol, 100 mM triethylammonium bicarbonate (TEAB, Sigma, T7408), pH 7.1 adjusted with phosphoric acid) before being applied to the S-Trap column. On-column digestion was carried out using mass spectrometry-grade Trypsin/Lys-C mix (Promega, V5073) at 37 °C in a humidified chamber. The eluted peptides were lyophilized, resuspended in 50 mM TEAB, and quantified using a quantitative fluorometric peptide assay (Pierce, 23290). The peptides were then labeled for 1 hour at room temperature with TMTpro18-plex reagent (Life Technologies, A44521 and A52048) at a 5:1 reagent-to-peptides mass ratio. Excess TMT reagent was quenched by adding hydroxylamine to a final concentration of 0.3%. The labeled samples were combined, lyophilized, desalted using Sep-Pak C18 Vac Cartridges (Waters, WAT023590), and fractionated with a reversed-phase fractionation kit (Pierce, 84868).

Mass spectrometry was performed using an Orbitrap Eclipse mass spectrometer equipped with a FAIMS Pro source, coupled to a Vanquish Neo nLC chromatography system using an EasySpray ES902 column (75 µm x 25 cm, 100 Ȧ) (Thermo Fisher Scientific). Mobile phases consisted of 0.1% formic acid (FA) in water (buffer A) and 0.1% FA in 80% acetonitrile (buffer B). Peptide fractions (1 µL) were injected and separated at 300 nL/min using a gradient of 3–25% B for 90 min, 25–40% B for 30 min, 40–95% B for 10 min, followed by 95% B for 6 min. The Orbitrap and FAIMS Pro were operated in positive ion mode with a 2100 V ionization voltage, a 300°C ion transfer tube temperature, and a 4.2 L/min carrier gas flow, using standard FAIMS resolution and compensation voltages of -45, –55, and –65 V. Full-scan MS spectra were acquired in centroid mode at a resolution of 120,000 (MS1 and MS2) over an m/z range of 400–1600, automatic maximum injection time, a custom AGC target (300% MS1, standard MS2), an isolation window of 0.7 m/z, an intensity threshold of 2.0 × 10⁴, charge state selection of 2-6, dynamic exclusion of 60 s, and HCD fragmentation with a collision energy of 55%. The RTS MS3 workflow was used. In RTS mode, the *Mus musculus* UniProt database was imported as the FASTA database, with trypsin specified as the enzyme for real-time spectral database searches. The search parameters included TMT 16-plex modifications on lysine and N-terminal amines (Δmass 304.2071), carbamidomethyl modification of cysteine (Δmass 57.0215), and a maximum of one variable methionine oxidation per peptide, with up to one missed cleavage. The RTS MS3 search was performed with a maximum search time of 40 ms. The MS3 scan was conducted using a mass range of 100–500 m/z, using an isolation window of 2 m/z for both MS and MS2. Peptide quantitation was carried out with a resolving power of 50,000 and a normalized collision energy of 55%. Additional parameters included a 200% normalized AGC target and automatic maximum injection time. Dynamic exclusion parameters were set at 1 within 60 s duration and ± 10 ppm mass window. All data were acquired using Xcalibur software (Thermo Fisher Scientific).

The PEAKS Studio 10.6 software package was used to analyze proteome data. Data extracted from the .raw files were first pre-processed with the following settings: scans were merged within a 10 ppm retention time window and a 10 ppm precursor m/z tolerance, including precursor mass and charge states (z = 2-8). Automatic centroiding, deisotoping, and deconvolution were then performed. Protein identification was performed using the *Mus musculus* UniProt database (accession UP000000589) with a parent mass error tolerance of 10 ppm, a fragment mass error tolerance of 0.5 Da, a retention time shift tolerance of 5.0 min, and semispecific trypsin specificity. TMTpro and carbamidomethylation (C) were set as a fixed modification, while variable modifications included oxidation (M), phosphorylation (P), and deamidation (NQ). One non-specific cleavage was allowed at either terminus, with up to three missed cleavages and a maximum of three variable post-translational modifications per peptide. False discovery rates (FDRs) were estimated using the decoy-fusion approach, with peptides considered confidently identified at an FDR ≤ 1% and a significance threshold of at least 20 (-10lgP). Protein quantification was based on the two most abundant peptide signals, which were used to calculate raw protein peak areas. Quantitative protein data were used to calculate fold-change differences between brain cell type-specific proteomes of aged and young mice. p-values were determined using a t-test with FDR correction by Benjamini and Hochberg.

### Immunoblotting

Lysates were mixed with 5X sample buffer (0.25 M Tris, 10% SDS, 25% glycerol, 25% β-Mercaptoethanol, and bromophenol blue) before being resolved by SDS-PAGE, followed by immunoblotting. For immunoblots in Figures 1, 2B and C, 4, S2 and S6, an equivalent of 0.1% and 12% of the whole tissue input and captured lysosomes were run, respectively. For immunoblots in Figures 5D and S14A, an equivalent of 0.1% and 5% of the whole tissue input and captured lysosomes were run, respectively. The following primary antibodies were used: Lamp1 (DSHB, 1D4B; RRID: AB_2134500), Lamp2 [ABL-93] (Abcam, ab25339; RRID: AB_470455), CtsD (R&D, AF1029; RRID: AB_2087094), CtsL (R&D, AF1515; RRID: AB_2087690), CtsB (CST, 31718S; RRID: AB_2687580), Citrate Synthase (CST, 14309S; RRID: AB_2665545), CALR (CST, 12238S; RRID: AB_2688013; and Abcam, ab92516; RRID: AB_10562796 for detection in rat samples), GAPDH (CST, 2118L; RRID: AB_561053), USO1 (Proteintech, 13509-1-AP; RRID: AB_2257094), ACOX1 (Abcam, ab184032; RRID: AB_2904240), NPC1 (Abcam, ab134113; RRID: AB_2734695), SDHA (CST, 11998S; RRID: AB_2750900), Vinculin (CST, 13901S; RRID: AB_2728768). HRP-conjugated secondary antibodies (anti-rat (RRID: AB_10694715) and anti-rabbit (RRID: AB_2099233)) were obtained from Cell Signaling Technology, and anti-goat HRP-labeled secondary antibody from Abcam (ab6741; RRID: AB_955424). HRP-conjugated mouse anti-rabbit conformation-specific antibody (CST, 5127S; RRID: AB_10892860) was used for immunoblotting of lysates obtained with the “tag-free” immunoisolation protocol. Immunoblots were visualized using film.

### Histology and Immunofluorescence

Brain tissue was fixed in 10% formalin, embedded in paraffin and sectioned at 5 μm. Antigen retrieval was performed using Borg Decloaker RTU solution (Biocare Medical, BD1000) in a pressurized Decloaking Chamber (Biocare Medical) for 3 min. Sections were blocked with 5% normal donkey serum and 0.3% Triton X-100 in PBS, followed by overnight incubation at 4°C with primary antibodies in 1% BSA and 0.3% Triton X-100. The next day, sections were incubated with secondary antibodies and Hoechst at room temperature before mounting with Vectashield Antifade Mounting Media (Vector Labs, H-1000). The following primary antibodies were used: HA-tag (CST, 2367S; RRID: AB_10691311), Iba1 (CST, 17198S; RRID: AB_2820254), NeuN (CST, 24307S; RRID: AB_2651140), GFAP (CST, 12389S; RRID: AB_2631098), Olig2 (CST, 65915S; RRID: AB_2936997). Alexa-488 and Alexa-568-conjugated secondary antibodies were obtained from Invitrogen. Fluorescent images were acquired on a Zeiss AxioVert 200M inverted microscope equipped with a Yokogawa CSU-22 spinning disk confocal scan head and a Hamamatsu Orca-ER cooled CCD camera. The MetaMorph acquisition software package was used to control image capture, ensuring consistent exposure times across slides.

### RNA extraction and RNA-seq

Total RNA was isolated from tissues using the miRNeasy Mini Kit (Qiagen, 217004) according to the manufacturer’s protocol, with modifications. Tissues were first cryogenically ground into a fine powder and then homogenized in Qiazol Lysis Reagent using the FastPrep-24 (MP Biomedical). The protocol was modified by incorporating 5PRIME Phase Lock Gel Heavy (QuantaBio, 232830) to facilitate easier phase separation.

RNA was further purified with 1.5X SPRI beads. RNA sequencing libraries were prepared with the NEBNext Ultra II Directional RNA Library Prep Kit for Illumina, using 250 ng of input RNA. Library preparation was automated with an SPT Mosquito HV liquid handler. The quality of indexed libraries was assessed using an Agilent Fragment Analyzer and qPCR. Sequencing was performed on a NovaSeq 6000, generating 50-nucleotide paired-end reads.

Paired-end reads (50x50) were mapped to the GRCm38/mm10 mouse genome with STAR v2.7.1a (*118*), using a STAR index created from gene annotation of canonical chromosomes from Ensembl Release 102 (*119*) merged with human TMEM192, with “--sjdbOverhang 49”. Genes were quantified with featureCounts v2.0.1 (*120*) using settings “-p -s 2”, and only protein-coding genes (as defined by Ensembl) were used for further analysis. Differential expression, assayed using DESeq2 v1.36.0 (*121*) with no log2FC shrinkage and “independentFiltering=FALSE”, was defined by a FDR < 1e-05, log2 fold change of at least 1, and mean normalized counts (“baseMean”) across the samples of interest higher than the median baseMean of all protein-coding genes.

### Statistical analyses

All polar metabolomics experiments were carried out at least two independent times, except for the analyses of female gut and uterus, as well as the rat samples and frozen samples obtained from the NIA, which were each performed once. Lipidomics analyses of mouse brain and heart samples were conducted two independent times, while analyses of liver and muscle were performed once. Proteomic analyses in Figure S11 were performed with one cohort for each brain cell type-specific analysis; full brain proteomics was conducted twice. Experimental samples with technical errors during extraction or processing were excluded from downstream analyses. Figure legends indicate the number of biological replicates (“n”). Statistical Quantitative data were visualized using Microsoft Excel, Python (via Google Colab), R (via RStudio Cloud), and GraphPad Prism 10. analyses were performed in Prism using two-tailed unpaired t-tests for pairwise comparisons or one-way ANOVA with post hoc tests for comparisons involving three or more groups, as specified. Multiple comparisons were corrected using the Benjamini-Hochberg method. Statistical significance was defined as p-value < 0.05. All displayed measurements represent independently generated samples or biological replicates. Immunoblot and immunofluorescence data are representative of biological replicates.

---

**Fig. S1.**
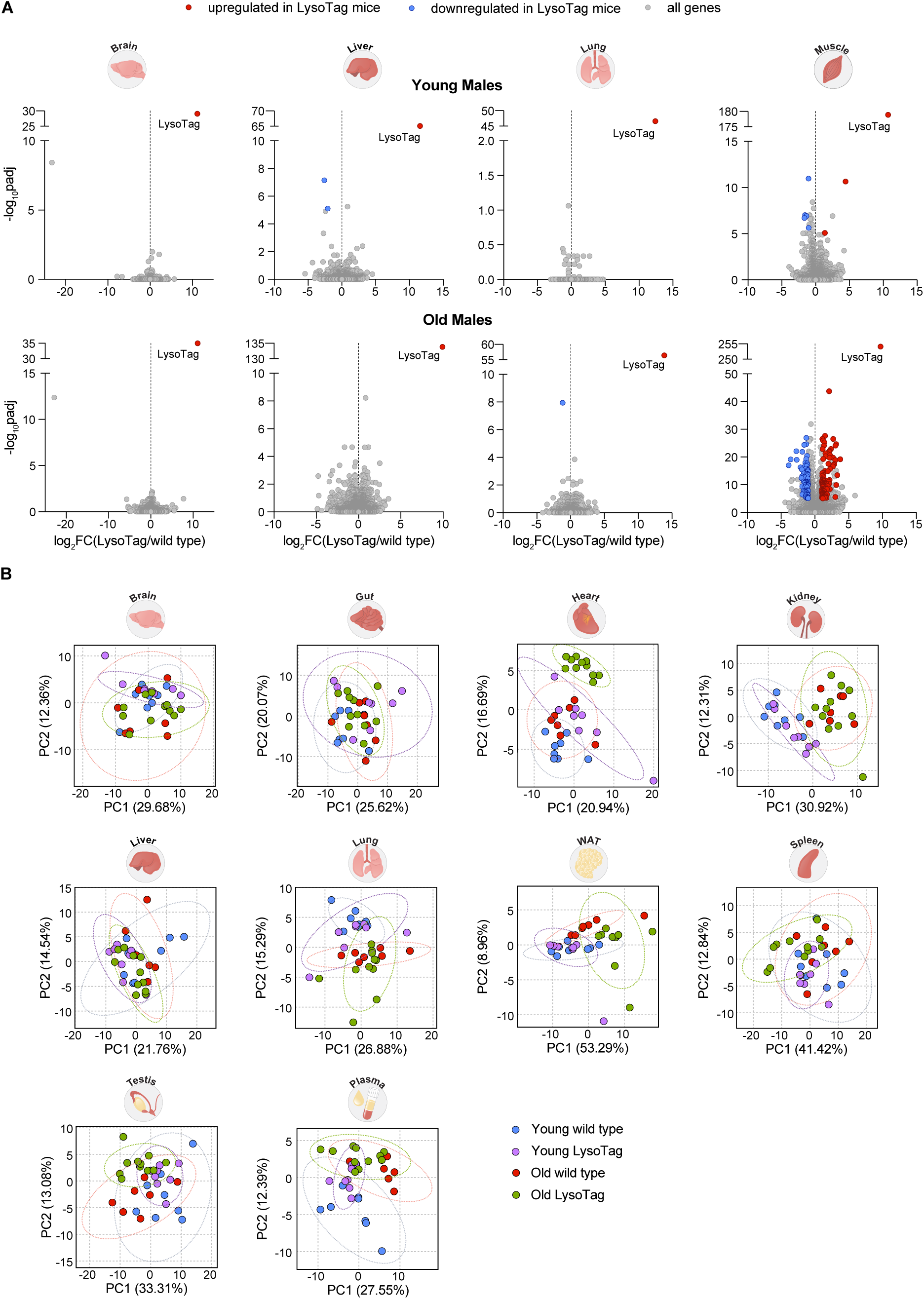
Comparison of whole tissue transcriptomes and metabolomes in LysoTag and wildtype mice. (A) Volcano plots depicting differentially expressed genes in the brain, liver, lung, and skeletal muscle (gastrocnemius) of young LysoTag (2 months old; n = 5) and wildtype (2 months old; n = 5) male mice (top); as well as old LysoTag (24 months old; n = 8) and wildtype (24 months old; n = 6) male mice (bottom). Genes upregulated in LysoTag mice are shown in red, while genes downregulated in LysoTag mice are shown in blue. (B) Principal component analysis (PCA) plots illustrating the effect of LysoTag expression and age on the targeted whole tissue and plasma polar metabolomes of male mice. The groups include young LysoTag (4-5 months old; n = 8), young wildtype (4-5 months old; n = 8), old LysoTag (24-26 months old; n = 9 to 11), and old wildtype (24-26 months old; n = 7). Ellipses indicating 95% confidence intervals are shown with dotted lines.

**Fig. S2.**
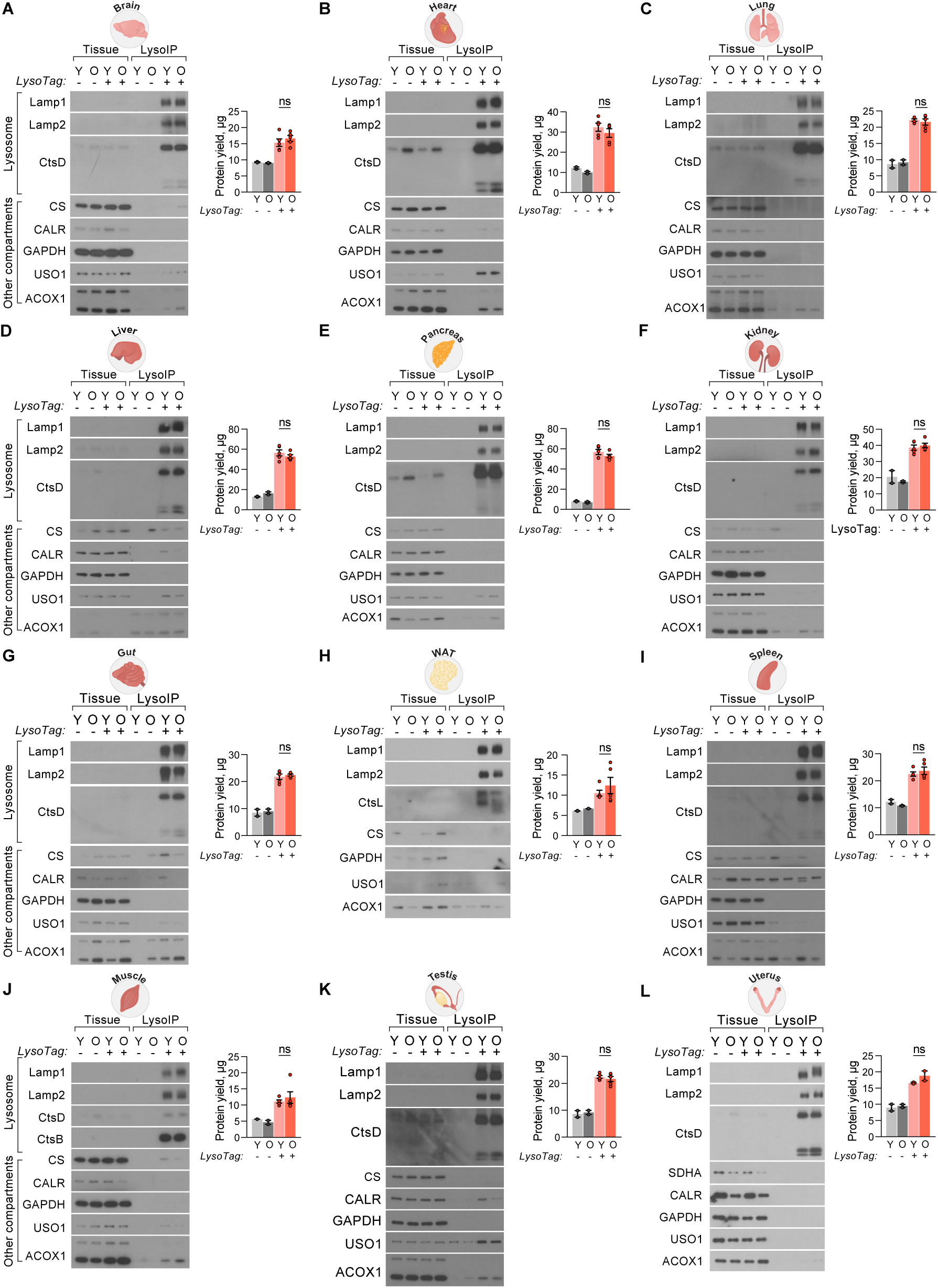
Lysosomes purified from tissues of young and aged LysoTag mice are of similar purity and yield. (A-L) Immunoblots of tissue lysates and lysosomal fractions from the indicated tissues of male mice (A-K), and female mice (L), probed for markers of lysosomes and other cellular compartments. A representative sample from each group is shown. Lamp1 – Lysosomal-associated membrane protein 1, marker of the lysosomal membrane; Lamp2 – Lysosomal-associated membrane protein 2, marker of the lysosomal membrane; CtsD – cathepsin D, marker of the lysosomal lumen; CtsL – cathepsin L, marker of the lysosomal lumen (probed in WAT), CtsB – cathepsin B, marker of the lysosomal lumen (probed in the muscle); CS – citrate synthase, mitochondrial marker; SDHA – succinate dehydrogenase complex flavoprotein subunit A, mitochondrial marker (probed in uterus); CALR – calreticulin, marker of the endoplasmic reticulum; GAPDH – Glyceraldehyde 3-phosphate dehydrogenase, cytoplasmic marker; USO1 – General vesicular transport factor p115, Golgi apparatus marker; ACOX1 – acyl-CoA oxidase 1, peroxisomal marker. Next to each immunoblot, the corresponding protein quantification is shown for young (n = 2) and old (n = 2) mock purification samples (LysoTag -), as well as young (n = 5) and old (n = 5) lysosomal fractions (LysoTag +). For the uterus samples, n = 2 mice per group. Note that in (A), the Lamp1, CtsD, and CS blots are <u>identical</u> to those in the abbreviated panel contained in Figure 1B.

**Fig. S3.**
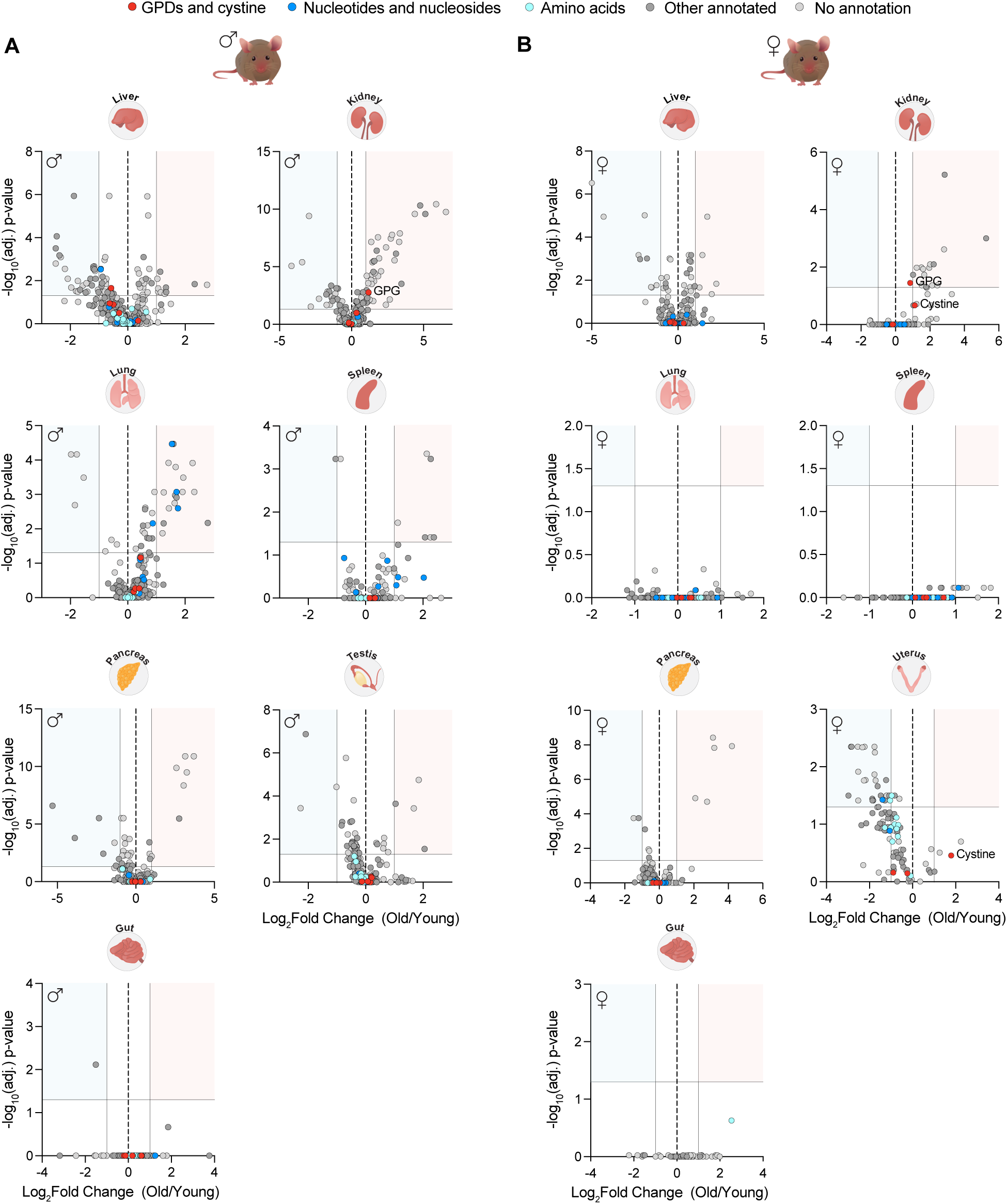
Comparison of lysosomal metabolomes from the tissues of young and old male and female LysoTag mice. (A-B) Volcano plots comparing untargeted lysosomal metabolomes from the indicated tissues of young and old mice. The gray horizontal line indicates an FDR-corrected p-value = 0.05, the vertical lines represent log_2_(Fold change) of -1 and 1. (A) Male cohort (young mice: 2-3 months old, n = 8 to 9; and old mice: 24-26 months old, n = 11). (B) Female cohort (young mice: 2-3 months old, n = 5 to 8; and old mice: 24-26 months old, n = 5 to 12).

**Fig. S4.**
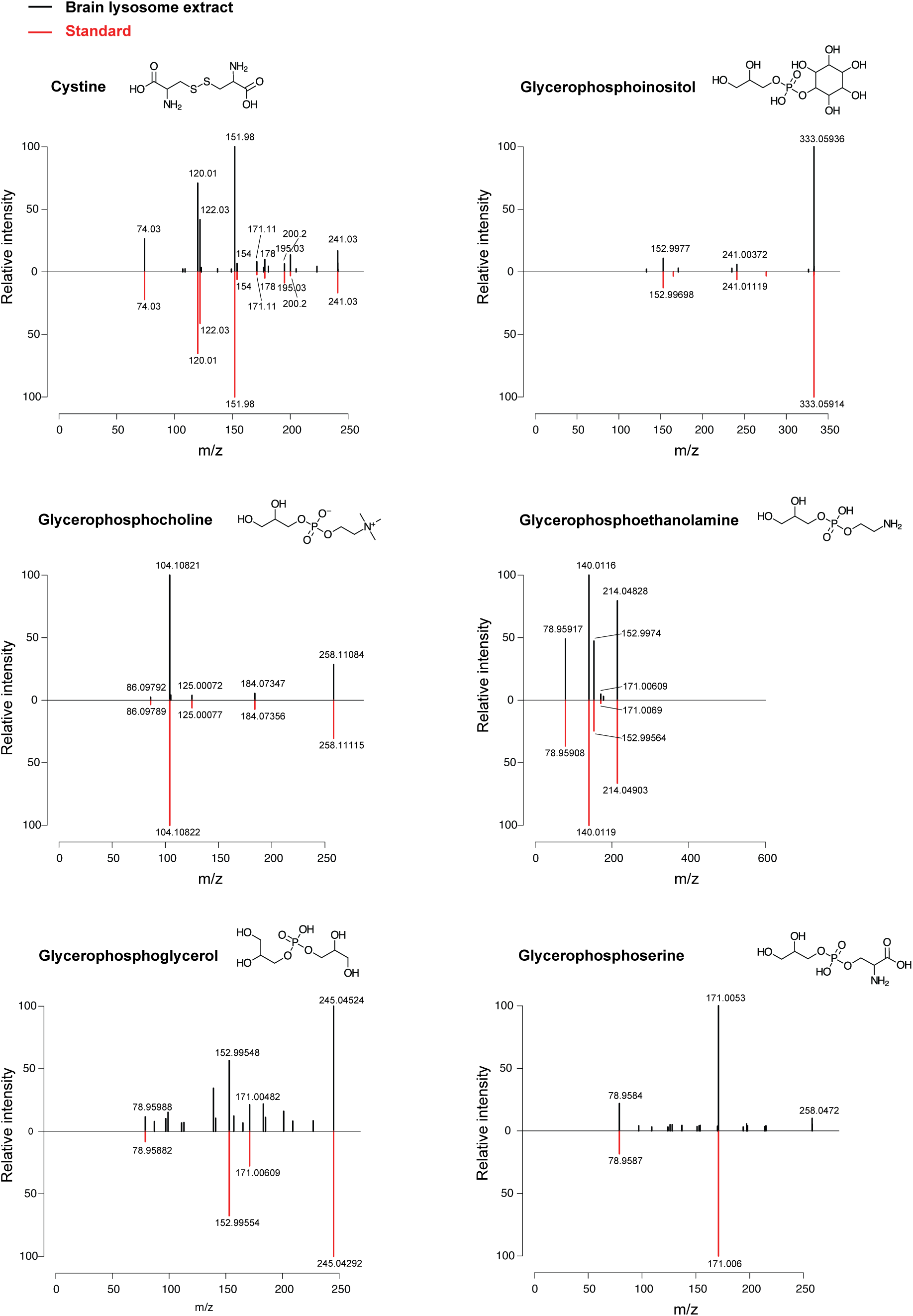
Validation of the presence of cystine and glycerophosphodiesters in metabolite extracts of brain lysosomes. Spectral mirror plots comparing cystine and glycerophosphodiesters in a representative lysosomal sample isolated from the brain of an old LysoTag mouse to its respective reference chemical standard.

**Fig. S5.**
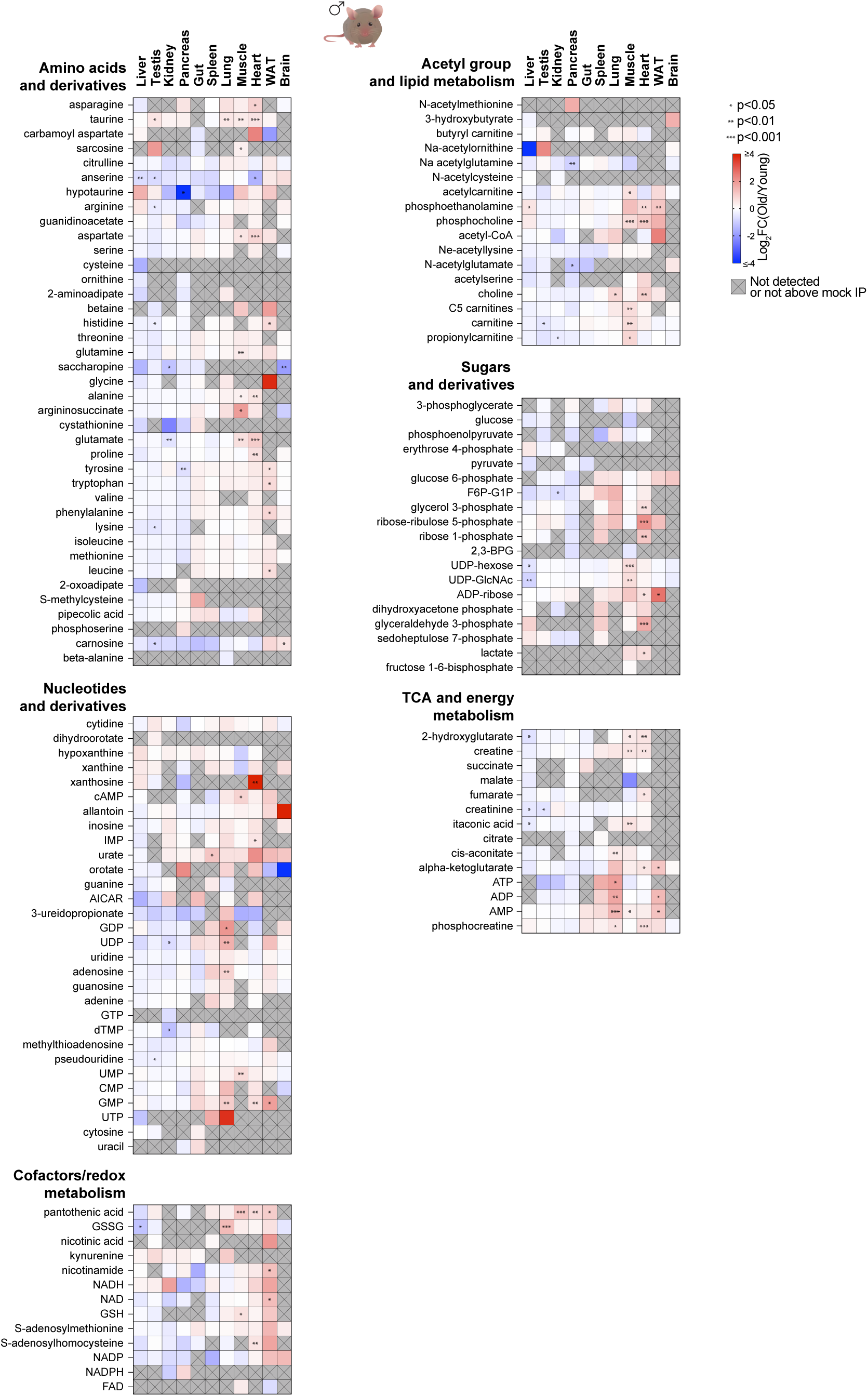
Targeted metabolite analyses in lysosomes isolated from young and aged LysoTag male mice. Heatmaps showing log_2_-fold changes of targeted metabolites in lysosomal fractions from aged relative to young mouse tissues (young male mice: 2-3 months old, n = 8 to 9; and old male mice: 24-26 months old, n = 11). Gray crossed tiles indicate metabolites that were not detected or did not meet the threshold of at least 2-fold enrichment in the lysosomal fraction relative to the mock purification. Asterisks denote FDR-corrected p-values (* p < 0.05, ** p < 0.01, *** p < 0.001).

**Fig. S6.**
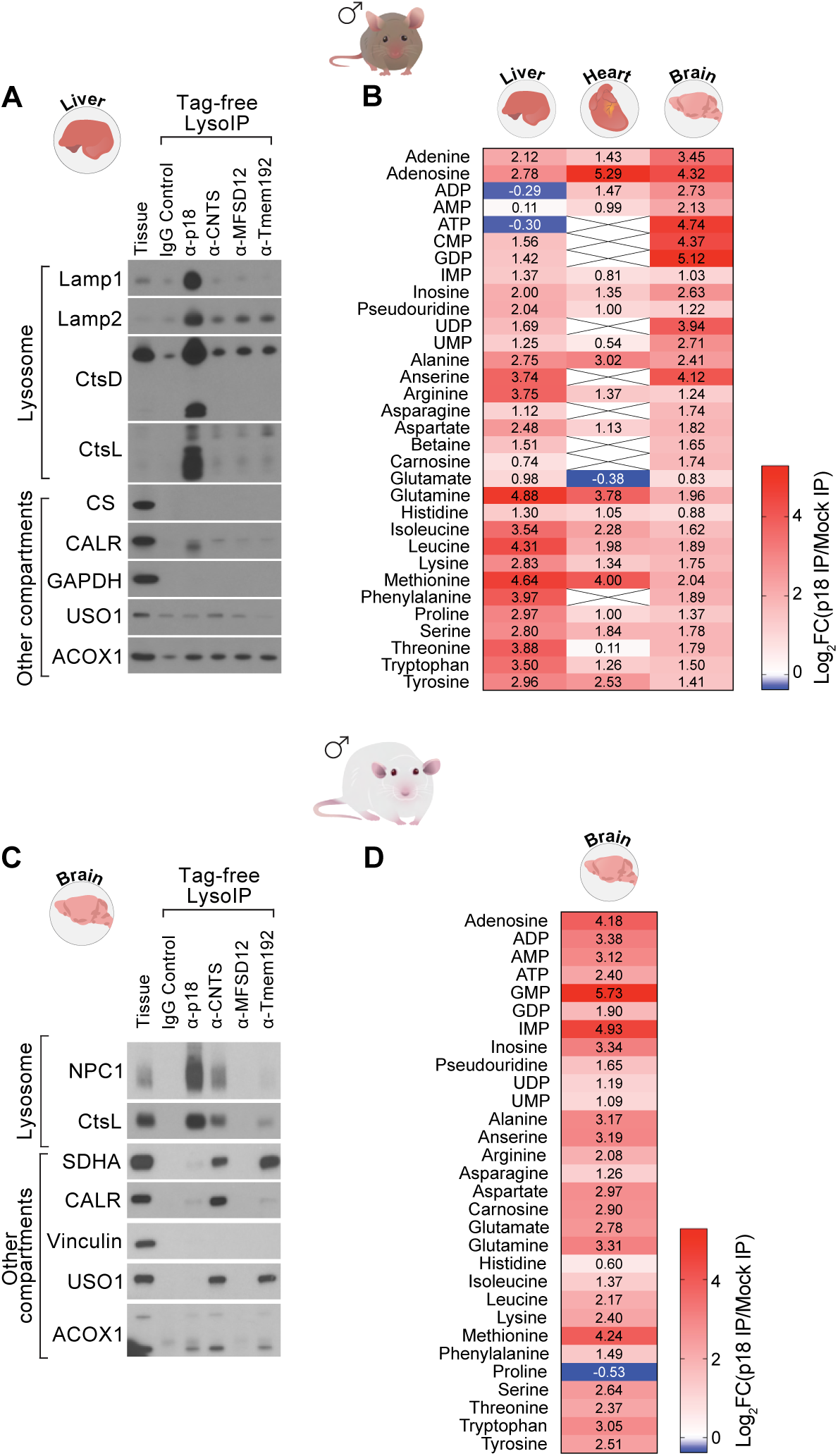
Development of “tag-free” lysosome immunoprecipitation from tissues of wildtype mice and rats. (A) Immunoblot analyses of mouse liver tissue lysates and lysosomal fractions using a panel of antibodies against lysosomal membrane proteins, along with rabbit IgG for a mock purification. Samples were probed for markers of lysosomes and other cellular compartments as in Figure S2. (B) Heatmap representation of log_2_-fold changes in a panel of targeted metabolites detected in lysosomal fractions obtained via anti-p18 immunoisolation from mouse liver (n = 3), heart (n = 4) and brain (n = 3), relative to mock purifications using isotype-specific IgG (n = 2 per tissue). Crossed tiles indicate metabolites that were not detected or not enriched above the mock purification. (C) Immunoblots of rat liver tissue lysates and lysosomal fractions obtained using a panel of antibodies against lysosomal membrane proteins, along with isotype-specific rabbit IgG for a mock purification. Samples were probed for markers of lysosomes and other cellular compartments: NPC1 – Lysosomal-associated membrane protein 1, marker of the lysosomal membrane; CtsL – cathepsin L, marker of the lysosomal lumen; SDHA – succinate dehydrogenase complex flavoprotein subunit A, mitochondrial marker, mitochondrial marker; CALR – calreticulin, marker of the endoplasmic reticulum; Vinculin, cytoplasmic marker; USO1 – General vesicular transport factor p115, Golgi apparatus marker. (D) Heatmap representation of log_2_-fold changes in a panel of targeted metabolites detected in lysosomal fractions obtained via anti-p18 immunoisolation from rat brain, relative to mock purifications with isotype specific IgG (n = 2).

**Fig. S7.**
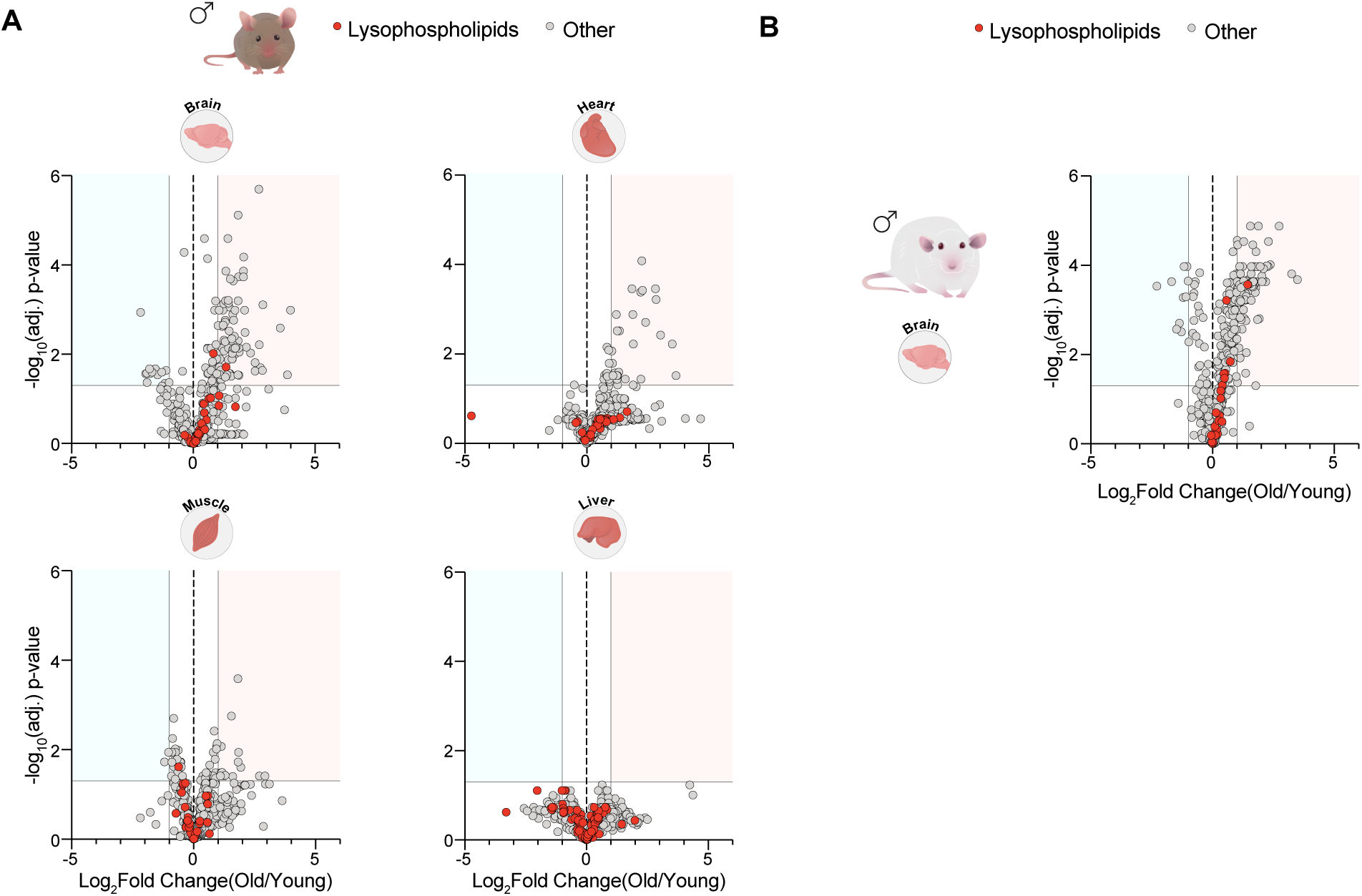
Lysosomes from aged wildtype mice and rats do not show globally elevated levels of lysophospholipids. (A) Volcano plots of untargeted lysosomal lipidomes from tissues of young (3 months old; n = 7) and old (24 months old; n = 7 to 9) wildtype male mice. The gray horizontal line indicates an FDR-corrected p-value = 0.05, the vertical lines represent log_2_(Fold change) of -1 and 1. Red points represent lysophospholipids. (B) Volcano plot of untargeted lysosomal lipidomes from the brains of young (4 months old; n = 8) and old (25 months old; n = 8) male rats. The gray horizontal line indicates an FDR-corrected p-value = 0.05, the vertical lines represent log_2_(Fold change) of -1 and 1. Red points represent lysophospholipids.

**Fig. S8.**
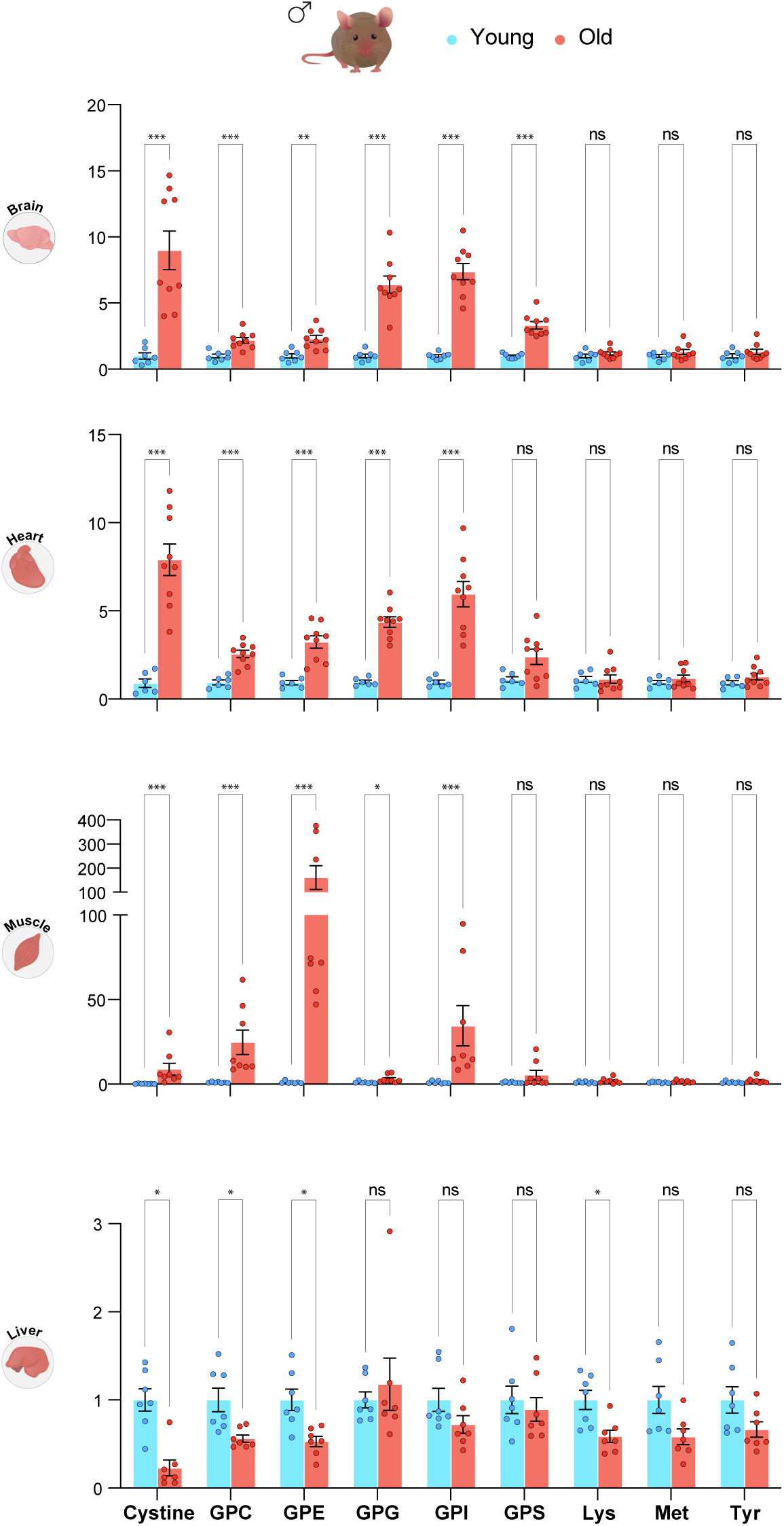
Verification of lysosomal glycerophosphodiesters and cystine levels in mice used for lysosomal lipidomics analyses. Relative levels of glycerophosphodiesters and cystine in the lysosome extracts generated for the untargeted lipidomics experiment, derived from tissues of young (3 months old; n = 6-7) and old (24 months old; n = 7 to 9) wildtype male mice. Asterisks denote FDR-corrected p-values derived from the Mann-Whitney test (* p < 0.05, ** p < 0.01, *** p < 0.001).

**Fig. S9.**
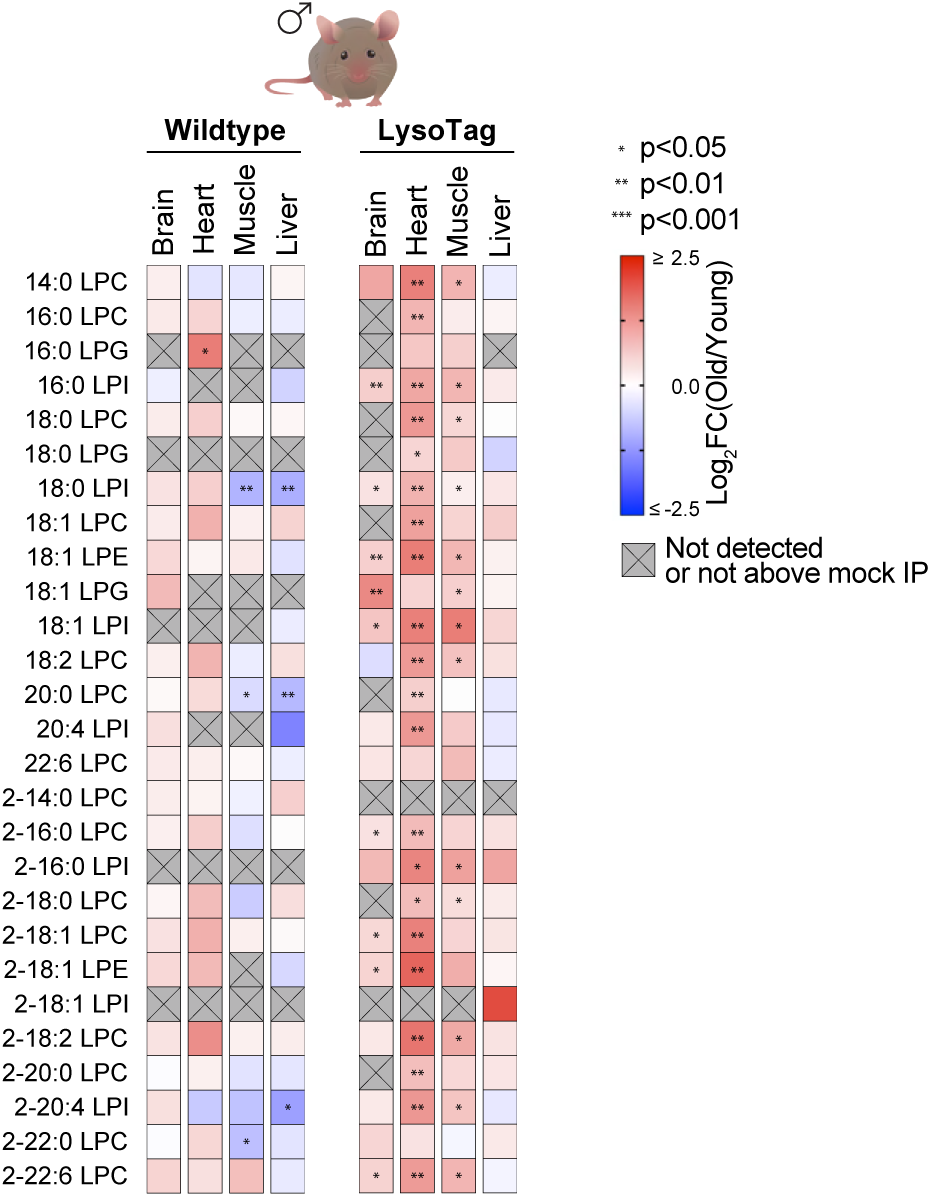
Lysophospholipid species are elevated in lysosomes from the brain, heart and muscle of the aged wildtype and LysoTag mice. Heatmaps showing log_2_-fold changes of targeted lysophospholipids in lysosomal fractions from aged relative to young wildtype (left, young male mice: 3 months old, n = 7; and old male mice: 24 months old, n = 7 to 9) and LysoTag mouse tissues (right, young male mice: 2-3 months old, n = 8; and old male mice: 24-26 months old, n = 8). Gray crossed tiles indicate metabolites that were not detected or did not meet the threshold of at least 2-fold enrichment in the lysosomal fraction relative to the mock purification. Asterisks denote FDR-corrected p-values (* p < 0.05, ** p < 0.01, *** p < 0.001).

**Fig. S10.**
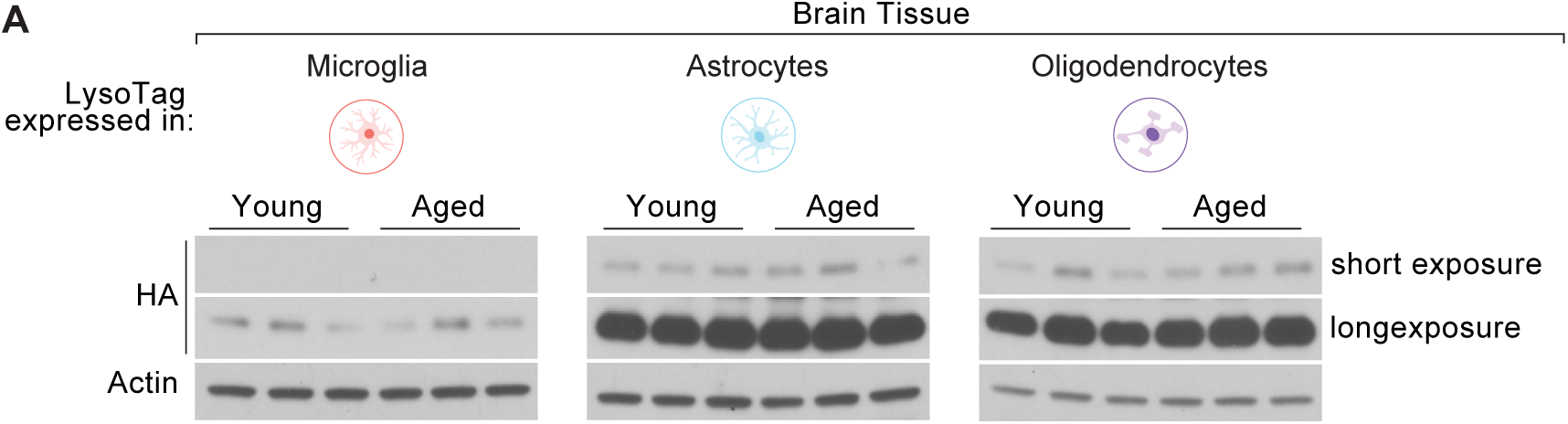
Tamoxifen-induced expression of the LysoTag in specific cell types of the brain. Immunoblots of whole-brain lysates from young (2-4 months old) and aged (24-25 months old) mice in which LysoTag expression was induced in specific cell types of the brain with tamoxifen. Blots were probed for HA and actin as a loading control (n=3 per group).

**Fig. S11.**
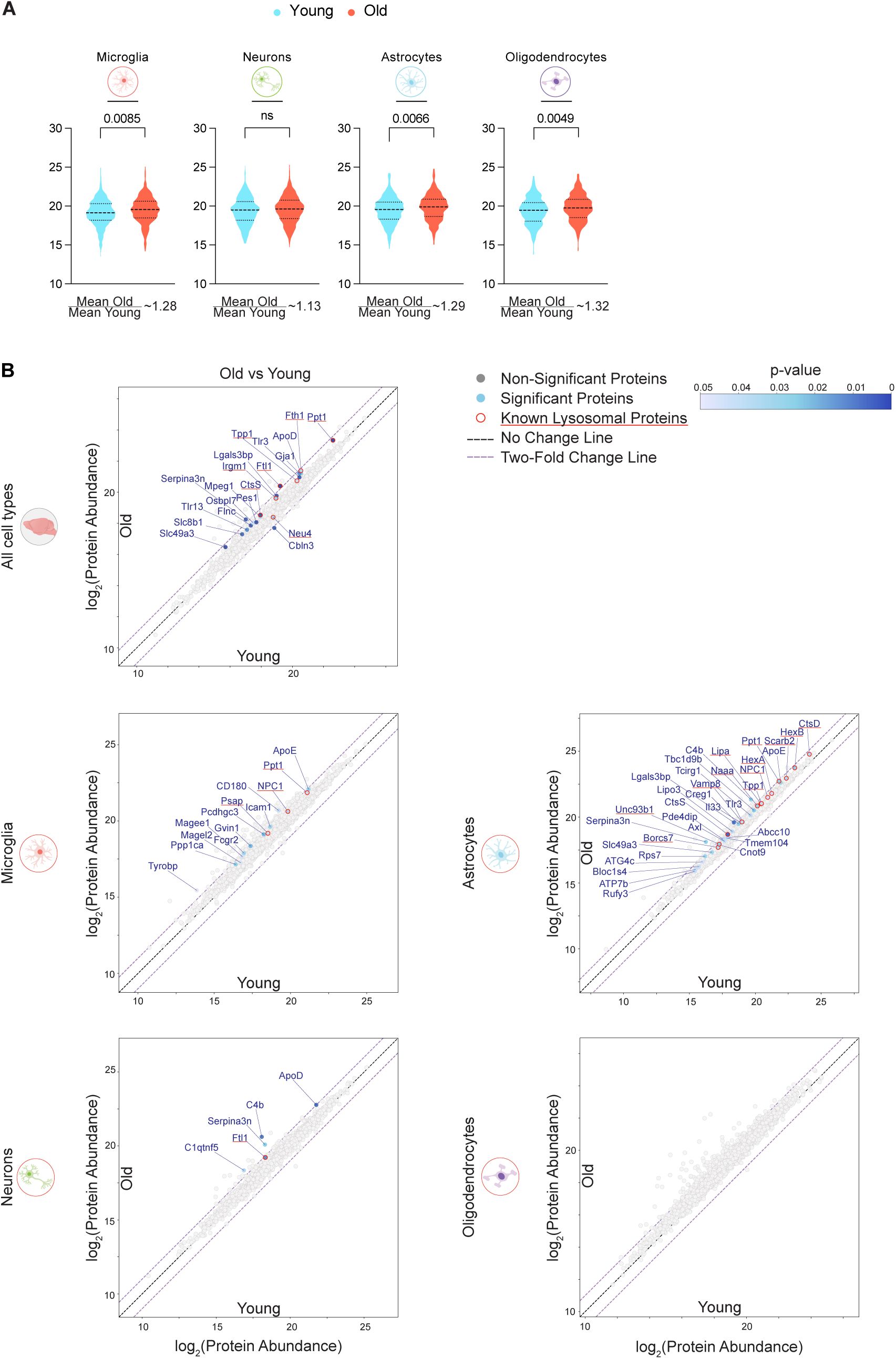
Comparison of lysosomal proteomes in the whole brain and specific brain cell types of young and aged mice. (A) Violin plots representing log_2_-transformed intensities of known lysosomal proteins in lysosomal fractions from young (3 months old; n = 6 to 8) and aged (23 – 26 months old; n = 5 to 8) mice expressing the LysoTag in specific brain cell types. Median, 25^th^ and 75^th^ percentiles are shown. The p-value indicates significance of the difference between the means of the populations. (B) Scatterplots comparing log_2_-transformed protein abundances in lysosomal fractions from the whole brain and specific brain cell types of young (3 months old; n = 6 to 8) and old mice (23 – 26 months old; n = 5 to 8). The x-axis represents young mice, and the y-axis represents old mice. Each dot corresponds to a protein, with the color scale indicating the FDR-corrected p-value. Proteins known to be lysosomal are highlighted with red outlines.

**Fig. S12.**
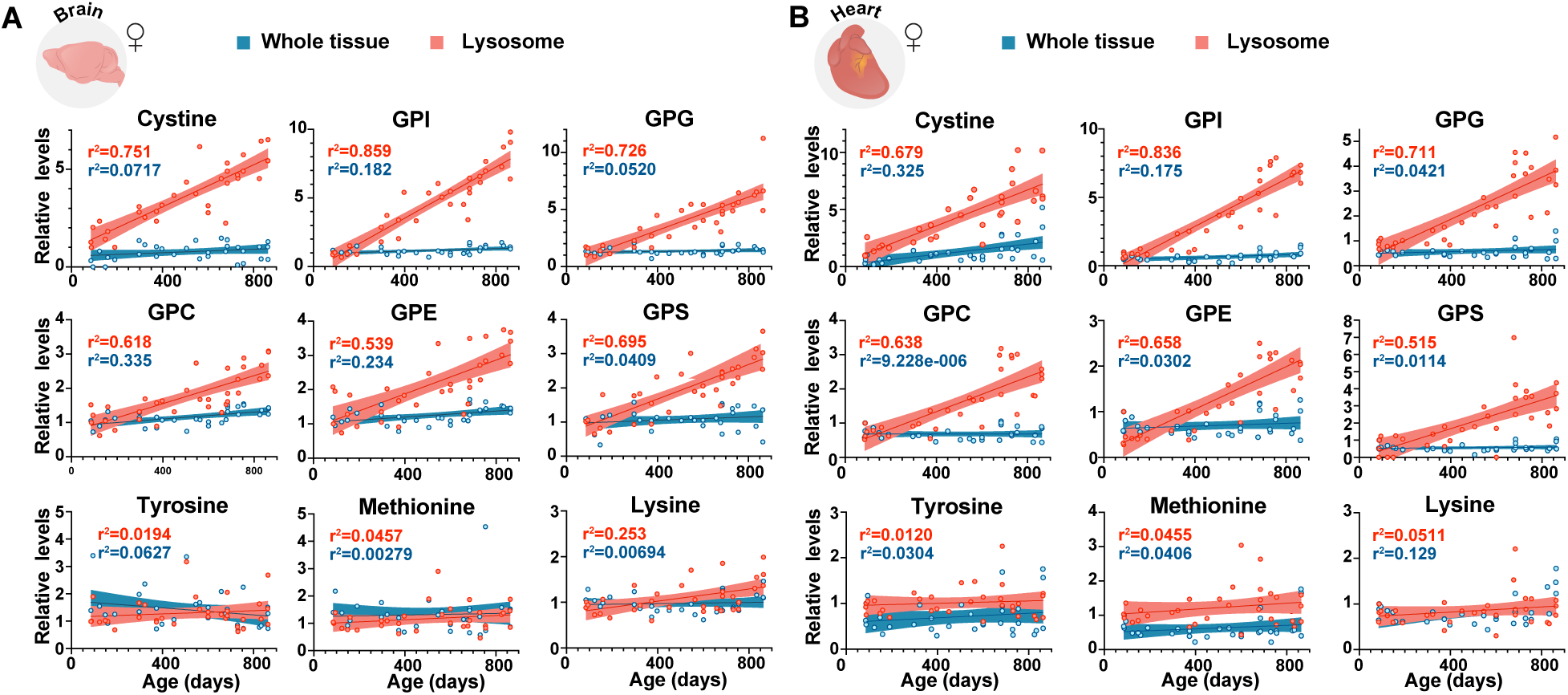
Lysosomal levels of cystine and glycerophosphodiesters increase with chronological age in wildtype female mice. (A) Levels of cystine, glycerophosphodiesters, and selected amino acids in brain lysosomes and whole brain tissue from female mice (n=33) ranging in age from 85 to 863 days. Shaded regions represent 95% confidence intervals around the regression line. (B) Levels of cystine, glycerophosphodiesters, and selected amino acids in heart lysosomes and whole heart tissue from female mice (n=32) ranging in age 85 to 863 days. Shaded regions represent 95% confidence intervals around the regression line.

**Fig. S13.**
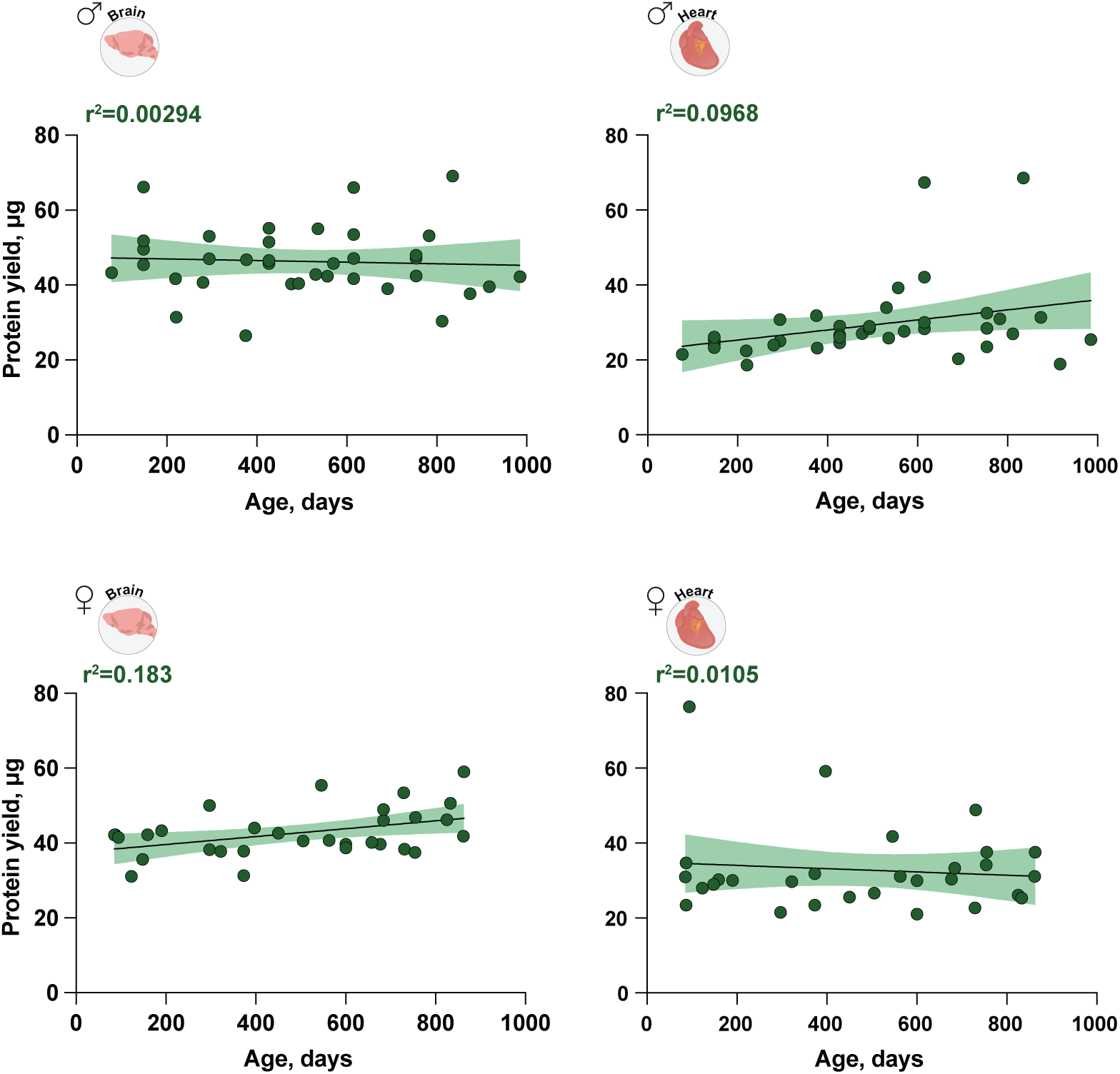
Quantification of protein yields in brain and heart lysosomes isolated from male and female mice across different ages. Shaded regions represent 95% confidence intervals around the regression line (n = 36 for male brain; n = 37 for male heart; n = 31 for female brain; n = 30 for female heart).

**Fig. S14.**
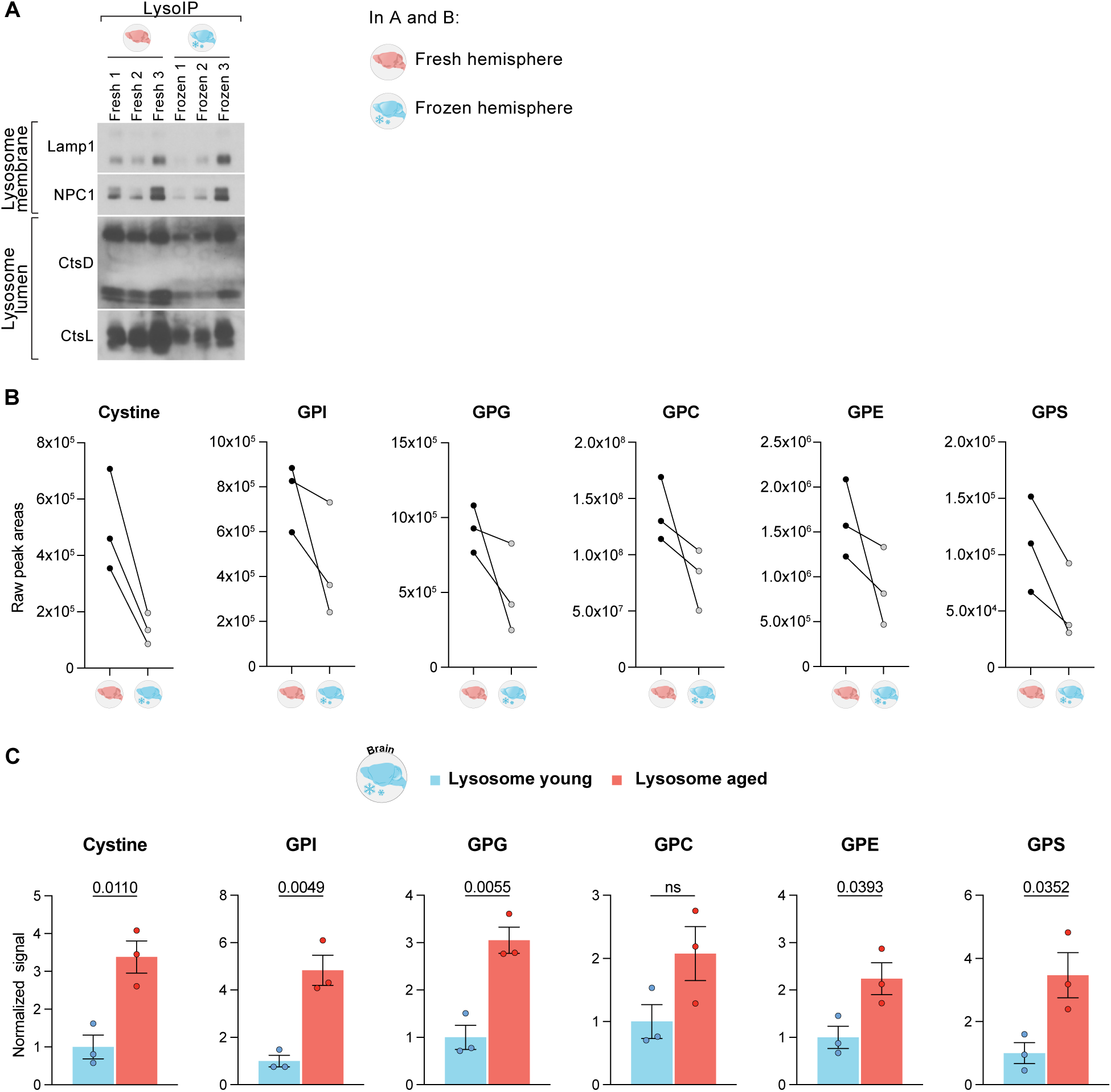
Comparison of lysosome extractions from fresh and frozen brain tissues. (A) Immunoblots of lysosomal fractions isolated from fresh and frozen brain tissue from 24-months old wildtype mice. Each brain was divided into the two hemispheres; one hemisphere was immediately used for “tag-free” lysosome isolation, while the other one was frozen in liquid nitrogen and stored at -80 °C for six weeks before lysosome isolation with the same “tag-free” method. Markers probed as in Figure S6. (B) Comparison of metabolite levels (cystine and glycerophosphodiesters) extracted from lysosomal fractions obtained in (A). (C) Comparison of metabolite levels (cystine and glycerophosphodiesters) in lysosomal fractions from young (3 months old; n=3) and aged (24-25 months old; n=3) wildtype mice using the “tag-free” lysosome purification from frozen brain tissues. Data are presented as mean ± SEM. p-values were determined using two-tailed t-test.

